# Extracellular vesicle-mediated secretion of chlorophyll biosynthetic intermediates in the cyanobacterium *Leptolyngbya boryana*

**DOI:** 10.1101/2024.03.29.587320

**Authors:** Kentaro Usui, Haruki Yamamoto, Hitoshi Mori, Yuichi Fujita

**Author notes:** Corresponding Author: K. Usui, Graduate School of Bioagricultural Sciences, Nagoya University, Furo-cho, Chikusa-ku, Nagoya 464-8601.

## Abstract

Extracellular vesicles (EVs) are derived from the outer membrane (OM) in Gram-negative bacteria and have diverse physiological functions such as toxin transport, extracellular signal transduction, nutrient acquisition, and horizontal gene transfer. EV-mediated secretion of monovinyl protochlorophyllide (MV-Pchlide), the chlorophyll *a* (Chl) biosynthetic intermediate, was previously reported in a mutant lacking dark-operative Pchlide reductase in the cyanobacterium *Leptolyngbya boryana*. This study showed a detailed characterization of EVs from the wild-type (WT) of *L. boryana* grown under photoautotrophic and dark heterotrophic conditions, focusing on the accumulation of Chl intermediates. WT *L. boryana* cells produce two types of EVs, low-density EVs (L-EVs) and high-density EVs (H-EVs), both under light and dark conditions. L-EVs and H-EVs showed distinct morphological features and protein compositions. L-EVs from cells grown under both light and dark conditions commonly contained carotenoids, myxol glycoside, and zeaxanthin, as major pigments. Based on the protein compositions of EVs and other cellular membrane fractions, L-EVs and H-EVs are probably derived from low-density OM and high-density OM interacting with cell walls, respectively. Fluorescence detection of pigments was applied to EVs, and the three Chl intermediates, protoporphyrin IX, demetallated Mg-protoporphyrin IX monomethyl ester, pheophorbide, and were commonly detected both L-EVs from light- and dark-grown cells, whereas L-EVs from dark-grown cells contained additional MV-Pchlide and MV-protopheophorbide. These Chl intermediates appear to transfer from the thylakoid membrane to L-EVs via an unknown transport system. Cyanobacterial EVs may play a novel function in alleviating the accumulation of Chl intermediates in cells.

## Introduction

Chlorophyll *a* (Chl) that absorbs the light energy from sunlight and initiates the photosynthetic electron transfer is an essential tetrapyrrole pigment for photosynthesis. Oxygenic photosynthetic organisms produce Chl from glutamate in a 15-step enzymatic reaction (Tanaka and Tanaka 2007; Masuda and Fujita 2008; Fujita and Yamakawa 2017). However, free Chl and its biosynthetic intermediates are potent, strong photosensitizers that generate reactive oxygen species harmful to cells upon light irradiation. Consequently, the Chl biosynthetic pathway is strictly regulated by a multilayer regulatory mechanism at transcriptional, translational, and post-translational levels to minimize the accumulation of free Chl and its intermediates (Czarnecki and Grimm, 2012; Wang et al. 2022). However, the full picture of the complex regulatory system remains still elusive. In particular, it is not fully understood how cyanobacteria, which do not have any Chl-degrading systems, unlike plants (Obata et al. 2019), efficiently remove free Chl and its intermediates generated in cells.

In the Chl biosynthetic pathway the first half-nine reactions from glutamate to protoporphyrin IX (PPN) are shared with the heme biosynthesis pathway. The latter six reactions specific in Chl biosynthesis (Mg branch; Fig. 1) begin from ATP-dependent insertion of Mg^2+^ ion into PPN, forming Mg-PPN by the action of Mg-chelatase. Subsequently, five successive reactions are followed: Mg-PPN methyltransferase, Mg-PPN monomethyl ester (MPE) cyclase, 8-vinyl reductase, protochlorophyllide (Pchlide) reductase, and Chl synthase, producing Chl. In the cyanobacterial Chl biosynthesis pathway two enzymes with different properties are involved in the four reactions: HemF and HemN for oxidative decarboxylation of coproporphyrinogen III (Goto et al. 2010); HemJ and HemY for protoporphyrinogen IX oxidation (Kobayashi et al. 2014); ChlA_I_ and ChlA_II_ for oxygen-dependent MPE cyclase (Minamizaki et al. 2008; Some species have another oxygen-independent MPE cyclase ChlE; Yamanashi et al. 2015); and light-dependent Pchlide reductase (LPOR) and dark-operative Pchlide reductase (DPOR) for the reduction of C17=C18 of Pchlide (Fujita 1996; Fujita et al. 1998; Yamazaki et al. 2006; Reinbothe et al. 2010). Utilizing analogous enzymes or isoforms with different properties and affinities for substrates, cyanobacteria can supply Chl without delay even in environments where light (high light, low light, and darkness) and oxygen levels (aerobic, hypoxic, and anaerobic) fluctuate (Fujita et al. 2015).

**Figure 1.**
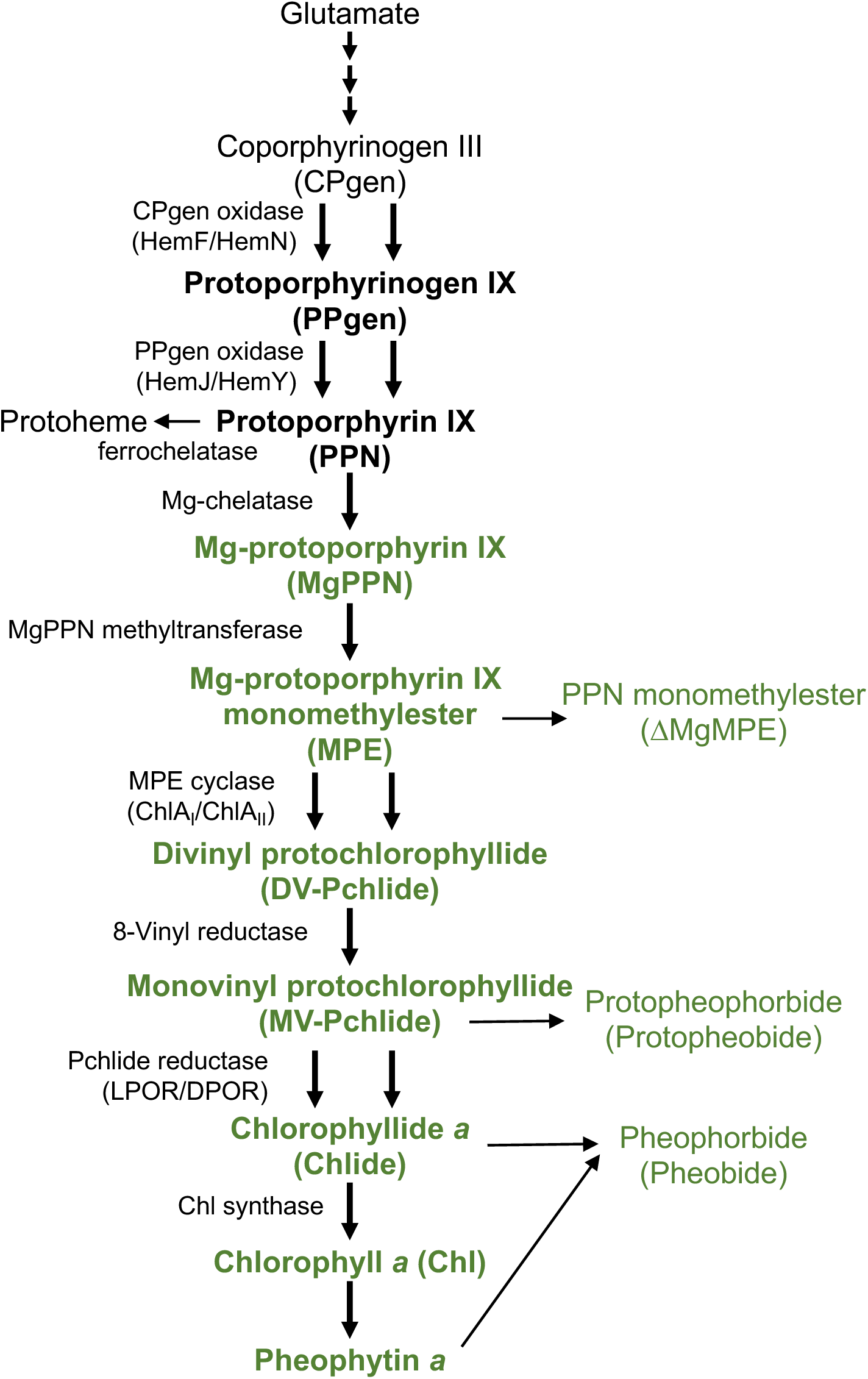
Chl biosynthesis pathway in cyanobacteria. Intermediates in the Mg branch of cyanobacterial Chl biosynthesis are shown in green. The steps at which two distinct enzymes differentially operate are shown with two arrows. Mg-depleted derivatives from intermediates of Mg-branch are shown by the thin arrows on the right.

In particular, the functional differentiation between DPOR and LPOR has been analyzed in detail in the cyanobacterium *Leptolyngbya boryana* based on the phenotype analysis of mutants lacking DPOR or LPOR. Under high light conditions, LPOR is almost solely responsible for Pchlide reduction (Fujita et al. 1998), whereas, under very low light or darkness, DPOR is the main enzyme for Pchlide reduction (Fujita et al. 1992; 1993; 1996; Kada et al. 2003). A mutant (ΔDPOR) lacking the DPOR subunit ChlL accumulates monovinyl (MV)-Pchlide in cells during dark heterotrophic growth, and furthermore secretes a large amount of MV-Pchlide, causing the culture medium to turn yellow. In the previous study, ΔDPOR secretes MV-Pchlide to the culture medium via extracellular vesicles (EVs; Usui et al. 2022). EV analysis revealed that ΔDPOR forms two EVs with different densities and that most of MV-Pchlide is secreted via high-density EVs (H-EVs). However, it is still unclear whether the EV-mediated secretion of MV-Pchlide is a special phenomenon caused only under unusual conditions where MV-Pchlide abnormally accumulates in cells, or whether WT *L. boryana* cells intrinsically produce EVs under normal growth conditions.

EVs are membranous vesicles derived from the plasma membrane (PM), and all bacteria can secrete EVs (Schwechheimer and Kuehn 2015). Gram-negative bacteria, which include cyanobacteria, are characterized by a three-layered cell membrane structure consisting of outer membrane (OM), cell wall (CW; peptidoglycan layer), and inner membrane (IM). EVs secreted by Gram-negative bacteria are derived from the OM (Kim et al. 2015). EVs contain various of biological substances, such as nucleic acids and proteins (Gonçalves et al. 2021). Furthermore, EVs have various physiological functions, including intercellular communication, biofilm formation, nutrient secretion and acquisition, toxin secretion, horizontal gene transfer, metabolite secretion, and response to environmental stress (Schwechheimer and Kuehn 2015). EV-mediated secretion of MV-Pchlide accumulated in cells can be regarded as a novel function of EVs (Usui et al. 2022).

In this study, EVs from WT *L. boryana* (dg5; Fujita et al. 1996; Hiraide et al. 2015) examined and characterized. WT also secrete two EVs with different densities regardless of photoautotrophic and dark heterotrophic conditions. Unlike EVs from ΔDPOR, WT secretes Chl intermediates mainly by low-density EVs. (L-EVs). Interestingly, Chl intermediates; PPN, demetallated MPE (ΔMgMPE), and pheophorbide (Pheobide), are commonly detected in both L-EVs from cells grown under light and dark culture conditions, and L-EVs from cells grown under dark heterotrophic conditions contained MV-Pchlide and the demetallated MV-Pchlide in addition to the three common pigments. These results suggest that *L. boryana* secretes Chl biosynthesis intermediates by L-EVs even under normal growth conditions and that EVs may play a novel regulatory role in Chl biosynthesis in cyanobacteria.

## Results

### EVs from WT grown under photoautotrophic and dark heterotrophic conditions

To examine whether the WT of *L. boryana* produces EVs under photoautotrophic (light) and dark heterotrophic (dark) conditions, the same method for isolation of EV fractions from ΔDPOR (Usui et al., 2022) was applied to prepare of the supernatants of WT cultures grown under light and dark conditions. The precipitates of the culture supernatants by ultracentrifugation were separated by stepwise sucrose density gradient centrifugation (Fig. 2). Orange bands appeared at the upper interfaces (0.6–1.6 M) from WT cells grown under both conditions (Fig. 2, upper panels). No clearly visible bands were observed at the lower interfaces (1.6–2.5 M). However, when the upper and lower interfaces were collected and they were concentrated by ultracentrifugation, not only the fractions from the upper bands gave rise to dark orange precipitates but also those from lower interfaces did small amounts of dark orange precipitates (Fig. 2, lower panels), suggesting that WT *L. boryana* grown under both light and dark conditions produced two EVs with different densities. These EVs fractions at the upper (0.6–1.6 M) and lower (1.6–2.5 M) interfaces were called L-EVs and H-EVs, respectively.

**Figures 2.**
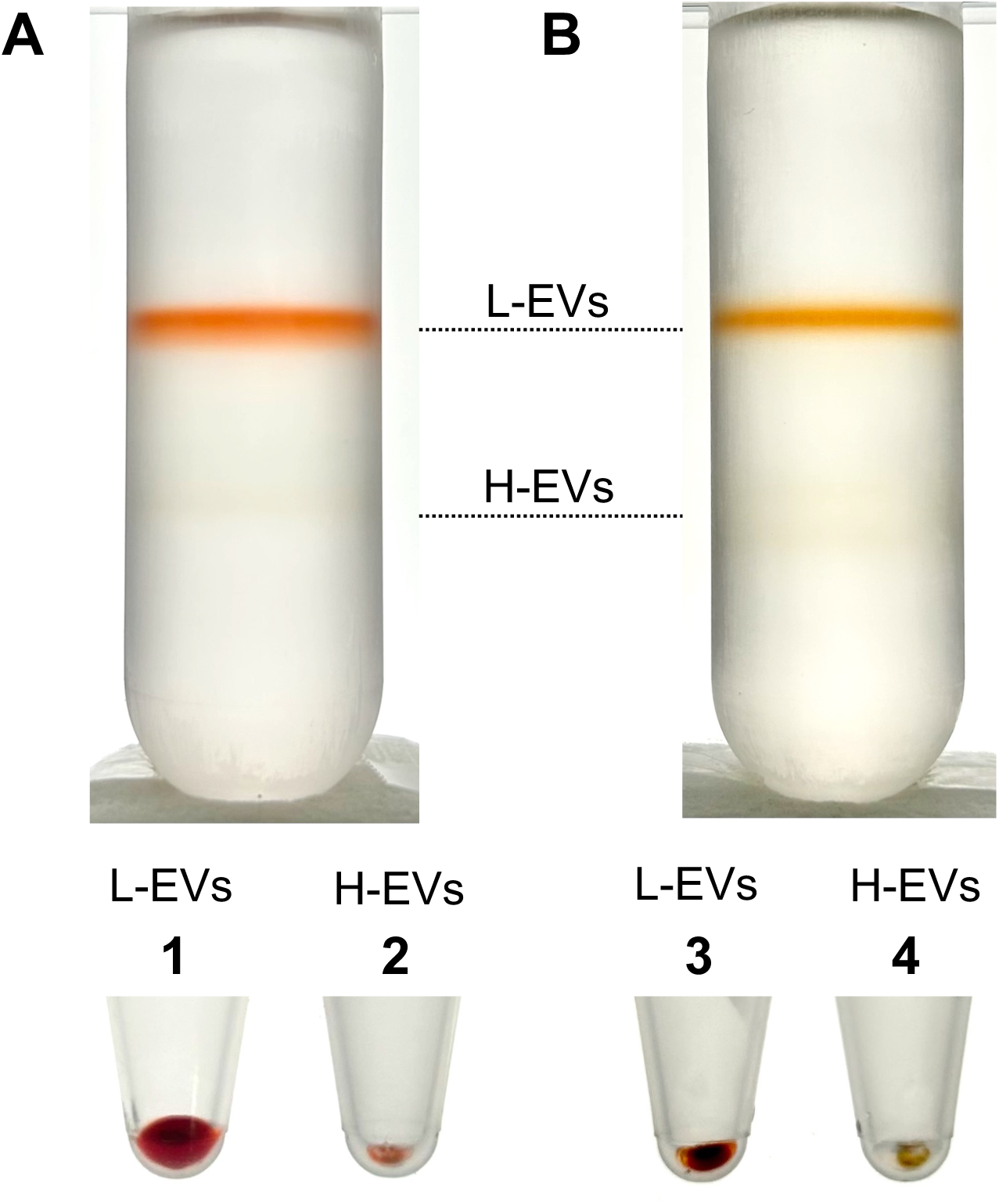
EVs isolated from WT grown under photoautotrophic and dark heterotrophic conditions. EV fractions from cells grown under photoautotrophic (**A**) and dark heterotrophic (**B**) conditions were isolated with stepwise sucrose density gradient centrifugation. EVs in the 0.6 M to 1.6 M and 1.6 M to 2.5 M interfaces were assigned as L-EVs and H-EVs, respectively. Precipitates of the EV fractions by ultracentrifugation are shown at the bottom. L-EVs (tubes 1 and 3) and H-EVs (tubes 2 and 4) were isolated from light-grown (tubes 1 and 2) and dark-grown (tubes 3 and 4) WT cells.

All four EV fractions were observed by transmission electron microscopy (TEM; Fig. 3). L-EVs and H-EVs exhibited respective common morphological characteristics irrespective of the growth conditions. L-EVs predominantly showed spherical structures (Fig. 3A and B), whereas H-EVs displayed bean-like unique structures with multiple vesicles interconnected (Fig. 3C and D), rather than a singular spherical shape.

**Figure 3.**
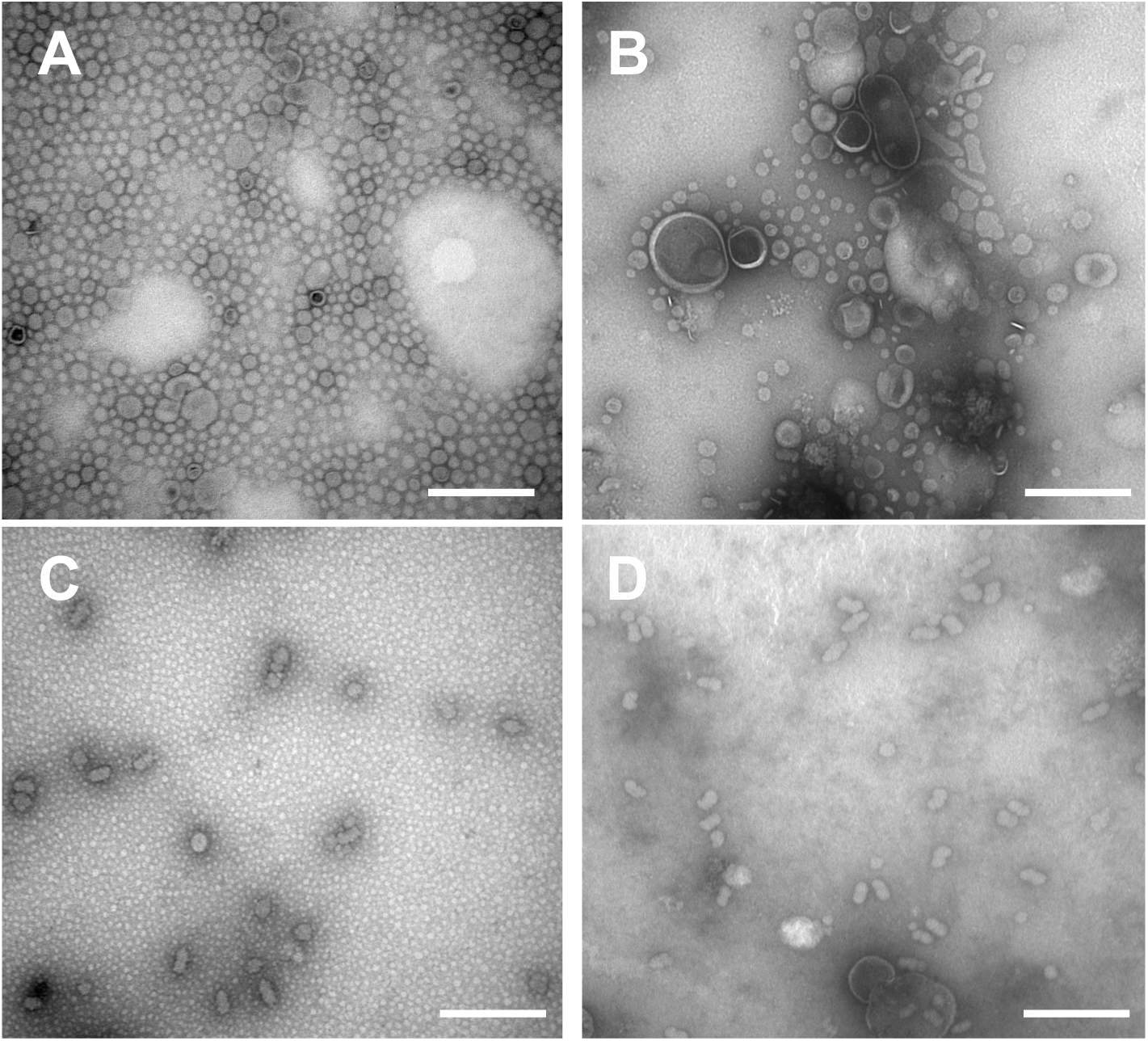
TEM observation of the four EV fractions. The EV fractions were observed by TEM with negative staining. L-EVs (**A** and **B**) and H-EVs (**C** and **D**) were isolated from cells grown under photoautotrophic (**A** and **C**) and dark heterotrophic (**B** and **D**) conditions. Bar, 300 nm.

Sodium dodecyl sulfate-polyacrylamide gel electrophoresis (SDS-PAGE) was performed to analyze the protein composition of these EV fractions (Fig. 4). The absence of thylakoid membrane (TM) or cell debris in all EVs fractions was confirmed by Western blot analysis using an antiserum against PsbA, photosystem II (PSII) reaction center protein D1. L-EVs and H-EVs obtained under both culture conditions showed distinct profiles. In both L-EV fractions from light and dark conditions, proteins with apparent molecular masses of ∼70 kDa and 50 kDa were predominant. In both H-EV fractions, multiple bands with apparent molecular masses of ∼50 kDa were observed as major proteins. The major proteins of EV fractions were identified by mass spectrometry (MS; Supplementary Table S1). The 70 and 50 kDa proteins in L-EV fractions were identified to be FG-GAP (phenyl-glycyl and glycyl-alanyl-prolyl consensus sequence; Springer 1997) repeat-containing protein (FGP; LBDG_07530: bands a and f) and porin isoforms (LBDG_25860 and LBDG_41910: band b; LBDG_41910: band g), respectively. The iron complex OM receptor proteins (LBDG_X2190 and LBDG_49870) were also detected in the 70 kDa band (band a) of L-EVs from the light-grown cells. The 50 kDa proteins in the H-EV fractions were identified as porin isoforms (LBDG_25860, LBDG_34630, LBDG_40860, and LBDG_41910: bands c–e; LBDG_25860, LBDG_40860, and LBDG_41910: bands h–j). The identification of the major proteins of EVs from WT was consistent with those of EVs from ΔDPOR in the previous study (Usui et al. 2022), indicating that under both culture conditions, WT and ΔDPOR produce two EVs that exhibit distinct densities and protein compositions.

**Figure 4.**
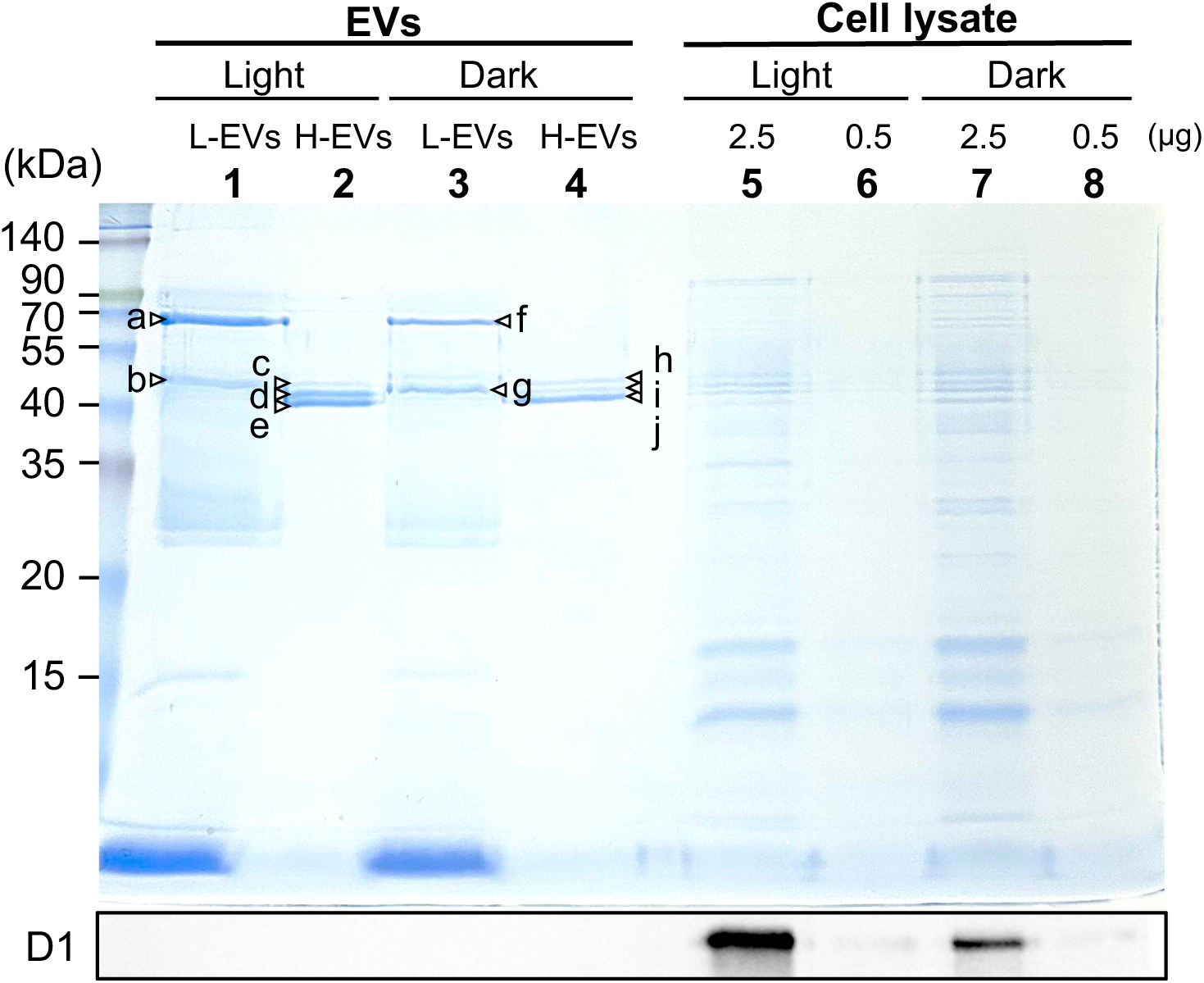
SDS-PAGE of L-EVs and H-EVs from WT cells grown under photoautotrophic and dark heterotrophic conditions. SDS-PAGE (CBB stain; top) of EVs fractions from WT cells grown under photoautotrophic (Light; L-EVs, lane 1; H-EVs, lane 2) and dark heterotrophic (Dark; L-EVs, lane 3; H-EVs, lane 4) conditions. An aliquot (2.5 µg protein) was loaded onto each lane. Protein bands shown with a to j were excised and analyzed by MS (Supplementary Table S1). Aliquots of CL were loaded onto lanes 5 to 8. As a marker protein for TM, D1 was detected with a specific anti-PsbA antiserum in the CL from WT cells grown under light (Light; 2.5 and 0.5 µg protein for lanes 5 and 6, respectively) and dark (Dark; 2.5 and 0.5 µg protein for lanes 7 and 8, respectively) conditions (Western blot analysis; bottom).

### Comparison of EVs to other membrane fractions

Cyanobacteria possess two types of membranes (TM and PM) and CW composed of peptidoglycan. Because cyanobacteria are Gram-negative bacteria, PM consists of two distinct membranes: OM and IM.

Three membrane fractions (TM, PM, and CW) were prepared from WT cells grown under light photoautotrophic (Fig. 5A) and dark heterotrophic (Fig. 5B) conditions using differential centrifugation and sucrose density gradient centrifugation (Murata and Omata 1988) (Supplementary Fig. S1A). PM fractions were precipitated by high-speed centrifugation from the supernatants of low-speed centrifugation of crude cell lysates (CL) and separated as orange bands at the interfaces of 0.6 to 1.6 M (Supplementary Fig. S1B and E). TM fractions were precipitated by low-speed centrifugation and separated by sucrose density gradient centrifugation to form dark green bands in the 1.6 M sucrose layers (Supplementary Fig. S1C and F). CW fractions were separated as orange bands at the interfaces of 1.6 to 2.5 M sucrose density gradient centrifugation after collecting orange bands that appeared just below the dark green TM bands (Supplementary Fig. S1D and G). To validate the characteristics of these fractions, protein compositions were compared using SDS-PAGE (Fig. 5) and Western blotting with an anti-D1 protein (PsbA) antiserum (Fig. 5, lower panels-). Protein profiles stained by Coomassie brilliant blue (CBB) revealed distinct protein compositions in all three fractions, and the three membrane fractions isolated from dark-grown cells showed nearly the same protein composition as that isolated from light-grown cells. The D1 protein of PSII was exclusively detected in CL and TM fractions, whereas it was absent in the PM and CW fractions. To identify the major proteins in PM and CW fractions, the main protein bands were excised from the gels and subjected to MS (Supplementary Tables S2 and S3).

**Figure 5.**
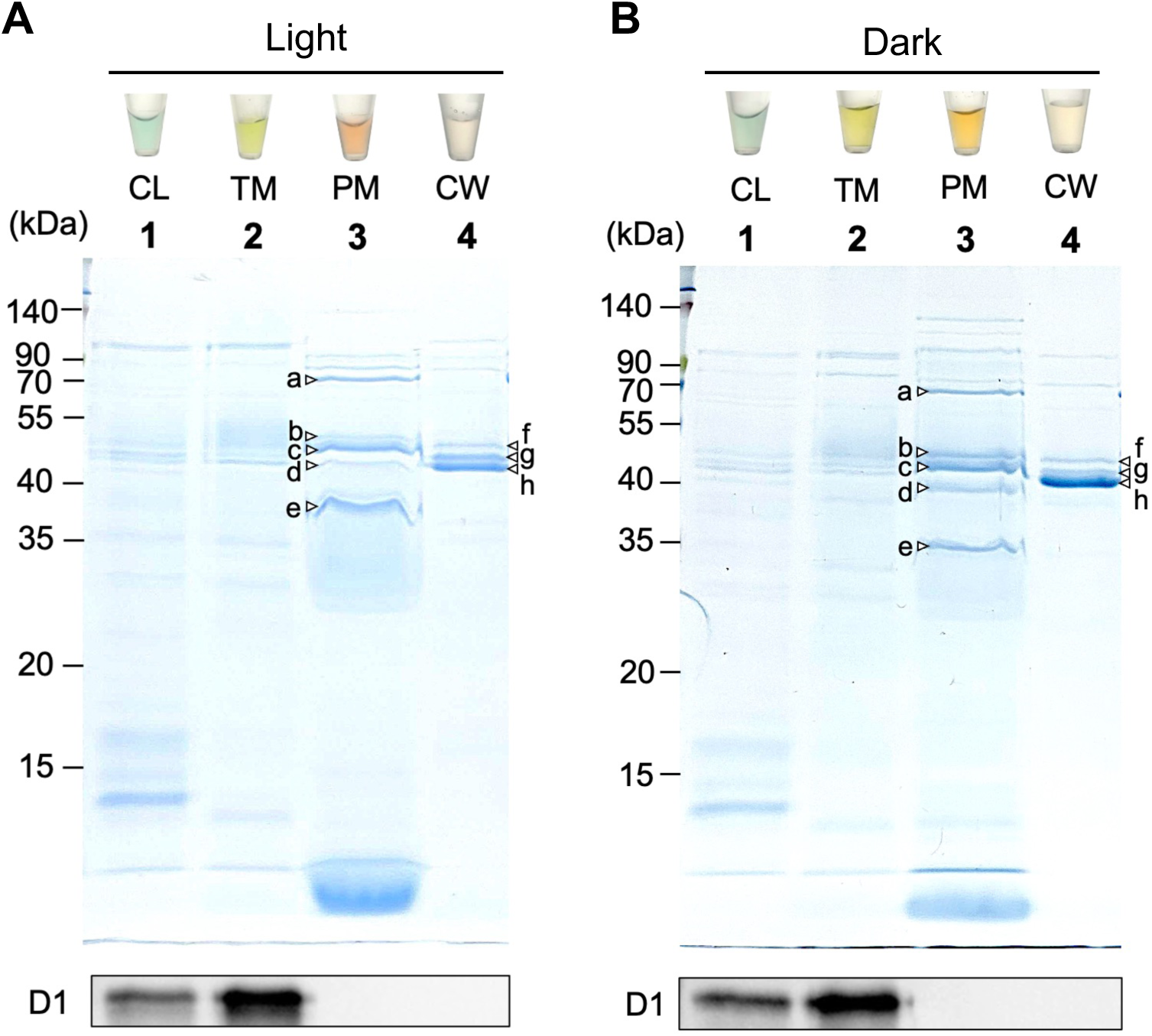
Membrane fractions from WT cells grown under photoautotrophic and dark heterotrophic conditions. SDS-PAGE (CBB stain; middle) and Western blot analyses (bottom) using the anti-PsbA antiserum for CL (lane 1), TM (lane 2), PM (lane 3), and CW (lane 4) isolated from WT cells grown under light (**A**) and dark (**B**) conditions, respectively. Top, suspension of the isolated fractions. Protein bands shown with a to h were excised and analyzed by MS (Supplementary Tables S2 and S3). An aliquot (2.5 µg protein) was loaded onto each lane.

In the PM fraction from light-grown cells, FGP (LBDG_07530: band a), iron complex OM receptor protein (LBDG_X2190: band a) and porin isoforms (LBDG_25860, LBDG_6050, and LBDG_40860: bands b and c) were identified. ABC transporter substrate binding proteins (LBDG_35200, LBDG_34900, LBDG_20220, and LBDG_5960: bands d and e), which were localized to IM (Norling et al. 1997), and phage shock protein A/IM 30 (LBDG_55170: band e), which was a homolog of the VIPP1 protein (56% similarity to VIPP1 from *Synechocystis* sp. PCC 6803 (PCC6803)) involved in TM biogenesis (Westphal et al. 2001), were also detected in the PM fraction. The protein composition suggested that the prepared PM fraction contained IM and OM. Comparing the protein composition between PM and L-EVs (Fig. 4), those of the two fractions were largely similar, although some were specific in PM, such as ABC transporter substrate binding proteins and the VIPP1 homolog, consistent with the idea that L-EVs are derived from OM of the PM fraction with lighter density.

The major proteins of the PM fraction from dark-grown cells (Fig. 5B), FGP (band a), porin (bands b and c), and VIPP1 homolog (band e), as well as that from light-grown cells, were also identified. In contrast, some proteins, such as starch synthases (band b) and NrtA (band d), were detected only in PM fractions from dark-grown cells. This difference would be interesting because it reflects the different growth modes between chemoheterotroph and photoautotroph, although this is not a proteomic analysis, as only the major proteins were excised for MS.

The SDS-PAGE profile of CW was highly similar to that of H-EVs. MS identified several porin isoforms (LBDG_6050, LBDG_17810, LBDG_25860, LBDG_40860, and LBDG_41910: bands f–h (light-grown cells; Fig. 5A), LBDG_25860, LBDG_40860, and LBDG_41910: bands f–h (dark-grown cells; Fig. 5B)) in the CW fraction. In PCC6803, porin isoforms (Slr1841, Slr1908, and Slr0042) were the major proteins of OM and also interacted with peptidoglycans (Kojima et al. 2016; Qiu et al. 2021). These porins possessed S-layer homology (SLH) domains interacting with peptidoglycans (Witwinowski et al. 2022). Based on these observations, the CW fraction predominantly comprised OM interacting with the CW showing higher density, and H-EVs were derived from OM interacting with the CW.

### Comparison of pigment compositions of EVs and other membrane fractions

To compare pigment compositions in the two EVs and other membrane fractions from photoautotrophically and dark heterotrophically grown cells, pigments were extracted from each fraction with 90% methanol and subjected to high-performance liquid chromatography (HPLC) analysis, and pigments were detected at an absorbance of 440 nm to assess overall pigment composition (Fig. 6). In the fractions from photoautotrophically grown cells, Chl (peak 3) was the predominant pigment in CL and TM, whereas almost no Chl peaks were detected in PM, CW, L-EVs, and H-EVs (Fig. 6A). In PM and L-EVs, carotenoids, specifically myxol glycoside (peak 1) and zeaxanthin (peak 2), were the common major pigments (Supplementary Fig. S2), whereas CW and H-EVs contained only trace levels of these carotenoids. The same carotenoid compositions of PM and L-EVs suggested that L-EVs were derived from PM, which was consistent with the idea that L-EVs are derived from OM with lighter density. This was also true between CW and H-EVs.

**Figure 6.**
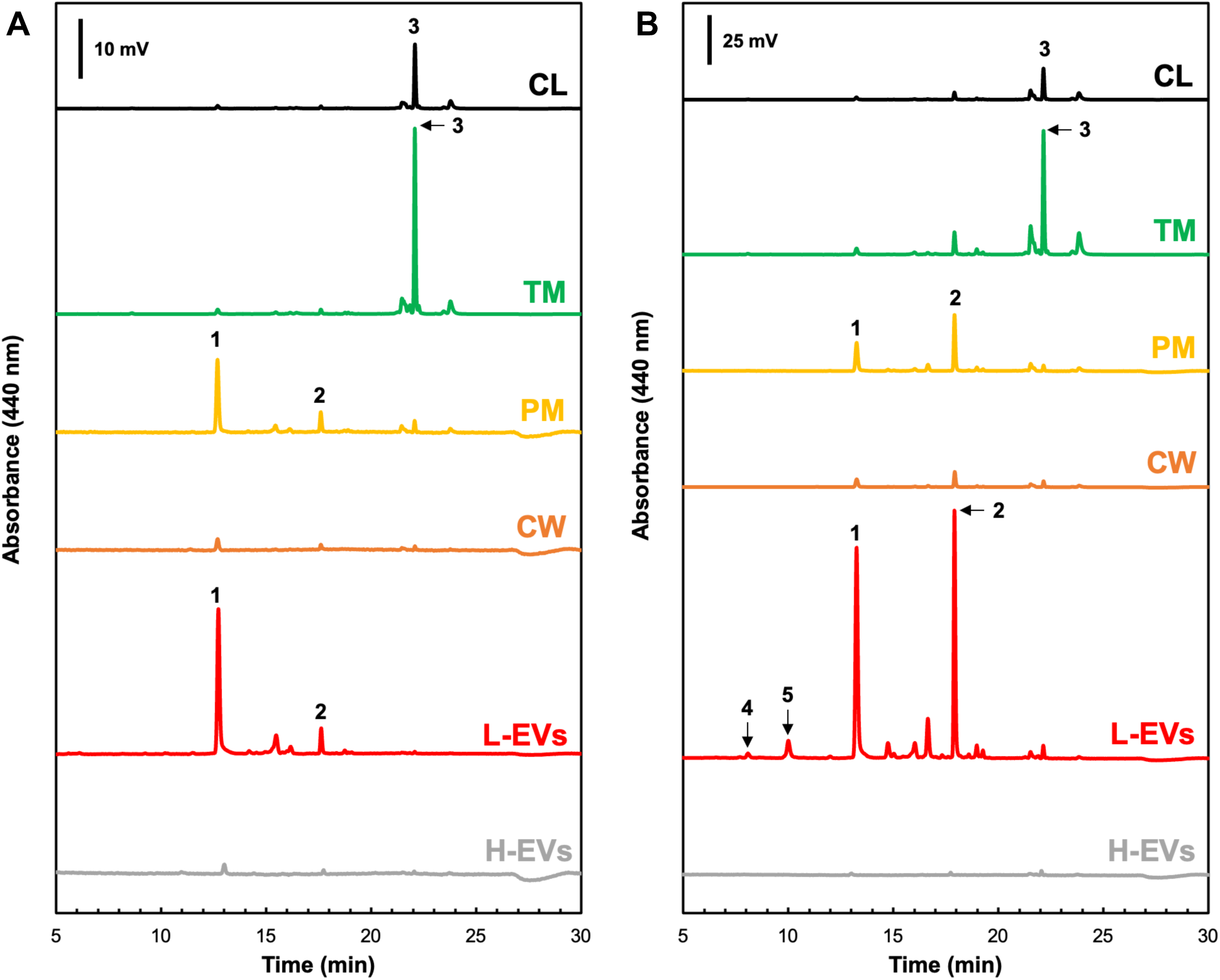
Pigment analysis of the L-EVs, H-EVs and other fractions from WT cells grown under photoautotrophic (**A**) and dark heterotrophic (**B**) conditions. Pigments in CL (black traces), TM, (green traces), PM (yellow traces), CW (orange traces), L-EVs (red traces), and H-EVs (gray traces) fractions were detected by absorbance at 440 nm. The protein concentration in each fraction was adjusted to 0.1 mg/ml, and an aliquot (20 µl) of each extracted pigment was injected. The peak assignment is as follows: 1, myxol glycoside; 2, zeaxanthin; 3, Chl; 4, MV-Pchlide; and 5, MV-Protopheobide. PDA absorption spectra of myxol glycoside (peak 1) and zeaxanthin (peak 2) are shown in Supplementary Fig. S2, and that of MV-Protopheobide (peak 5) is shown in Supplementary Fig. S3.

EVs and membrane fractions from cells grown under dark-heterotrophic conditions showed slightly different pigment compositions (Fig. 6B). Chl was the major pigment in CL and TM, and myxol glycoside and zeaxanthin were the major pigments, which had the same features as in PM and L-EVs from the light-grown cells. However, the amounts of carotenoids were much higher than those in PM and L-EVs from the light-grown cells. The ratio of myxol glycoside to zeaxanthin was almost equal in PM and L-EVs from the light-grown cells. In addition, small amounts of MV-Pchlide and MV-protopheophorbide (MV-Protopheobide) were detected in L-EVs (Fig. 6B, peaks 4 and 5; Supplementary Fig. S3).

Subsequently, to explore the presence of Chl biosynthetic intermediates in the membrane fractions in higher sensitivity, fluorescence spectroscopic detection at two different wavelengths was conducted in the HPLC analysis (dashed line, λ_em_/λ_ex_ 660/430 nm; solid line, λ_em_/λ_ex_ 630/400 nm; Fig. 7). In this HPLC condition, Chl intermediates before the attachment of phytol were eluted from 5 to 15 min (Aoki et al. 2014). In the fractions from light-grown cells (Fig. 7A), Chlide (peak 1) and a Chlide derivative (peak *) were detected as the major Chl intermediates in TM (Supplementary Figs. S4 and S5). Pheobide (peak 3; Supplementary Fig. S3) and Chlide were mainly detected in PM (Supplementary Fig. S4). Interestingly, PPN (peak 2) and ΔMgMPE (peak 4) were detected in L-EVs in addition to Pheobide (Supplementary Fig. S6). PPN and ΔMgMPE were specific to L-EVs and not detected in other membrane fractions.

**Figure 7.**
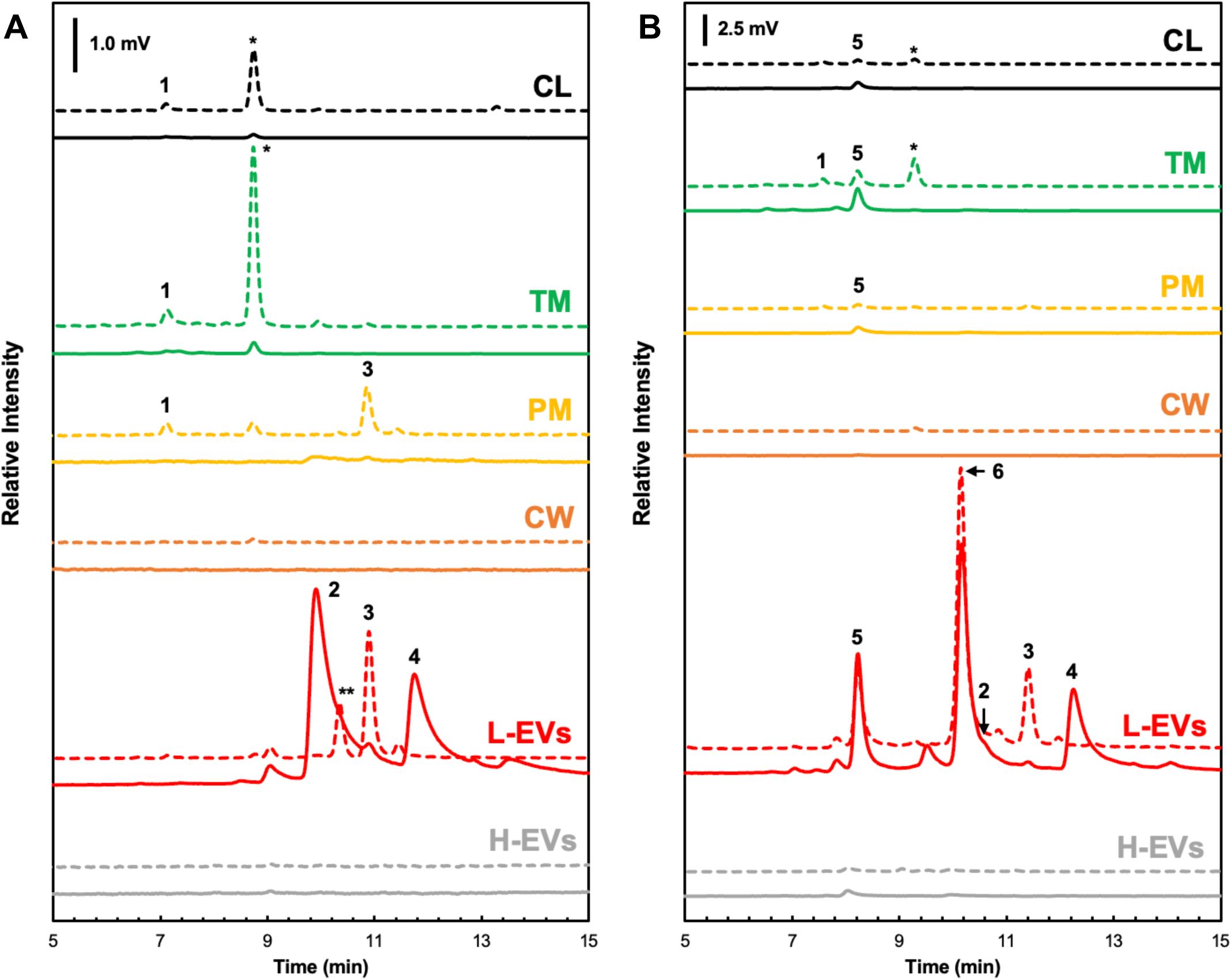
Chl intermediates in the EVs and other fractions from cells grown under photoautotrophic conditions (**A**) and dark heterotrophic (**B**) conditions. Chl intermediates are detected by fluorescence emission (dashed lines, λ_em_/λ_ex_ 660/430 nm; solid lines, λ_em_/λ_ex_630/400 nm) in CL (black traces), TM (green traces), PM (yellow traces), CW (orange traces), L-EVs (red traces), and H-EVs (gray traces). The protein concentration of each fraction was adjusted to 0.1 mg/ml, and an aliquot (20 µl) of each extracted pigment was injected. The peak assignment is as follows; 1, Chlide; 2, PPN; 3, Pheobide; 4, ΔMgMPE; 5, MV-Pchlide; and 6, MV-Protopheobide. Peaks * and ** were not identified but pigment of peak * would be a Chlide derivative (Supplementary Fig. S5).

In TM from dark-grown cells (Fig. 7B), MV-Pchlide (peak 5) was detected. A very small amount of MV-Pchlide was also detected in PM. In L-EVs, MV-Pchlide, MV-Protopheobide (peak 6), PPN, Pheobide, and ΔMgMPE were detected. The three pigments, PPN, ΔMgMPE, and Pheobide, were common intermediates in L-EVs from light- and dark-grown cells, whereas MV-Pchlide and MV-Protopheobide were specific in L-EVs from dark-grown cells. No Chl intermediates were detected in H-EVs, regardless of light or dark conditions.

HPLC analysis was conducted for CL and L-EV fractions using fluorescence wavelengths (λ_em_/λ_ex_ 600/417 nm) to detect specifically Mg-PPN and MPE (Supplementary Fig. S7). MPE was detected as a minor peak in CL fractions from dark-grown cells and L-EVs from light-grown cells. However, Mg-PPN (eluted at 7 min; Aoki et al. 2014) was not detected in any fractions.

Taken together, the results suggested that (1) the major pigments of EVs were two carotenoids (myxol glycoside and zeaxanthin); (2) the contents and ratio of the two carotenoids differed in L-EVs from cells grown under light and dark conditions; (3) based on the protein compositions, L-EVs and H-EVs were derived from low-density OM and high-density OM interacting with CW, respectively; and (4) Chl intermediates accumulate in L-EVs. The three intermediates (PPN, ΔMgMPE, and Pheobide) were commonly detected in both L-EVs, whereas L-EVs from dark-grown cells contained additional intermediates (MV-Pchlide and MV-Protopheobide).

## Discussion

In this study we found that the filamentous cyanobacterium WT *L. boryana* secreted two EVs (L-EVs and H-EVs) of different densities under photoautotrophic and dark heterotrophic culuture conditions. L-EVs and H-EVs commonly contain porin isoforms, but exhibit distinct protein and carotenoid compositions. The protein and pigment compositions suggested that L-EVs and H-EVs are derived from low-density OM and high-density OM interacting with CW, respectively. In addition, L-EVs contain Chl biosynthetic intermediates and their derivatives, depending on the culture conditions.

### EV formation

Based on the results, a model for EV formation in WT *L. boryana* was proposed (Fig. 8A). The similarity between L-EVs and PM in density, pigment composition, and protein composition was consistent with that of L-EVs derived from PM. The PM fraction contained both proteins localized in IM and OM (Supplementary Tables S2 and S3), whereas L-EVs contained mainly proteins localized in OM (Supplementary Table S1) and the content of FGP was higher than PM (Figs. 4 and 5). This feature suggested that L-EVs originate from the OM region where shows less interaction with CW than the other regions and FGP is localized predominantly (Fig. 8A). This model was consistent with that L-EVs have a lower content of porin isoforms compared to the CW fraction, that contains OM interacting with the CW (Figs. 4 and 5).

**Figure 8.**
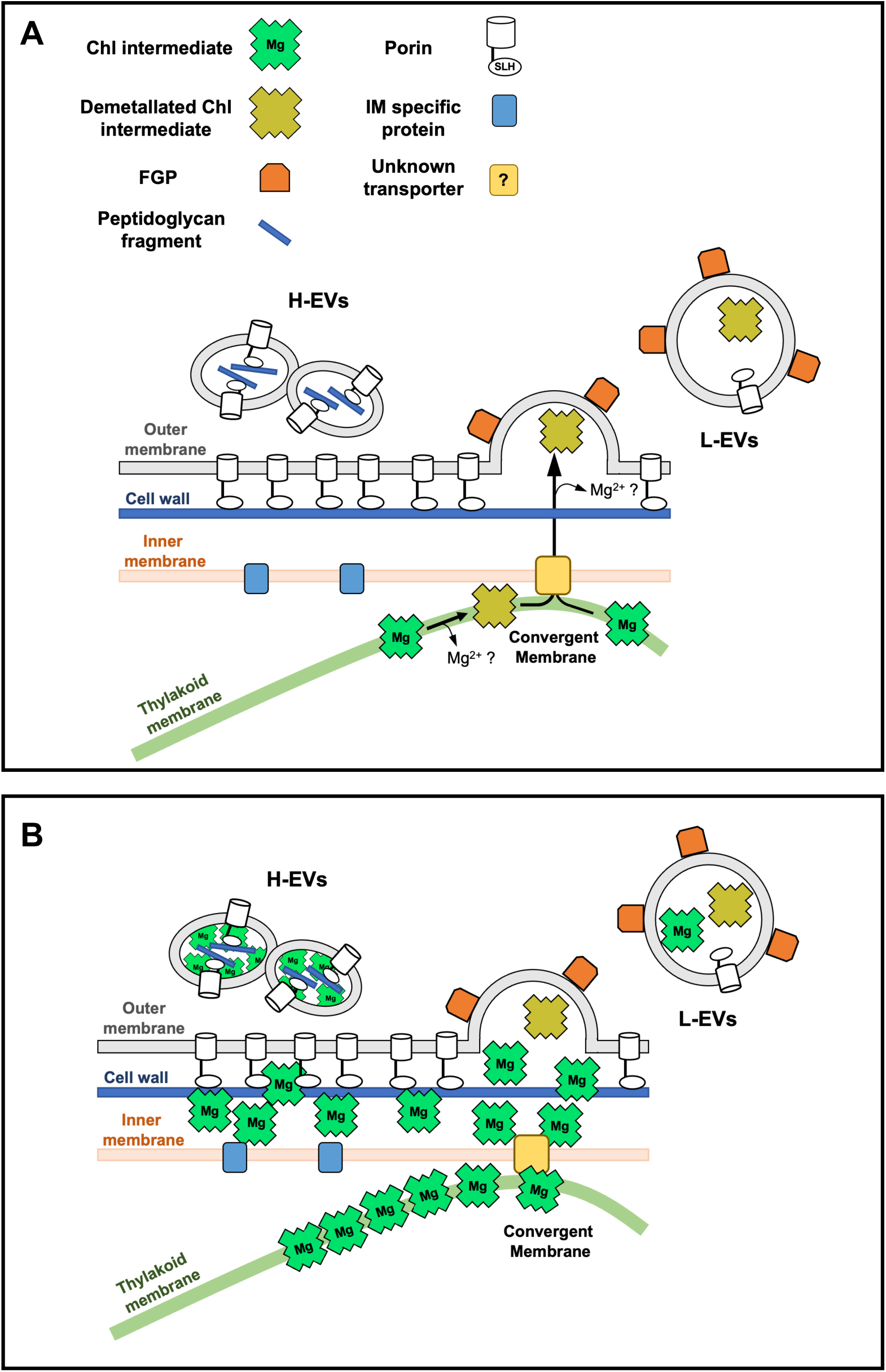
Models for the formation of two EVs in *L. boryana* cells. **A.** WT *L. boryana* secretes two EVs under light and dark conditions: L-EVs, whose major protein and pigments are FGP and carotenoids, respectively, and H-EVs, whose major protein is porin and contains much less amounts of pigments than L-EVs. In WT, free Chl intermediates accumulated intracellularly are loaded into only L-EVs and secreted out from the cells. L-EV-mediated secretion could alleviate the accumulation of Chl intermediates within the cell. **B.** *L. boryana* ΔDPOR also secretes two EVs. During the dark heterotrophic growth of ΔDPOR, a large amount of MV-Pchlide accumulates in the cells. Such abnormally accumulated MV-Pchlide is mainly secreted via H-EVs (Usui et al. 2022). A differential loading system for L-EVs and H-EVs is still unknown.

H-EVs differed significantly from L-EVs in density, protein composition and shape. H-EVs had a unique bean-like shape, and some vesicles linked each other (Figs. 2 and 3C and D), different from the spherical shape of L-EVs. The pigment and protein compositions of H-EVs were more similar to the CW fraction than the PM fraction (Figs. 4 and 5). Porin isoforms are the major proteins in H-EVs. Porin contains the N-terminal SLH domain that interacts specifically with peptidoglycans (Witwinowski et al. 2022; Kojima et al., 2016). These features suggested that H-EVs form from the OM regions where porin isoforms are more densely present and interact extensively with the CW (Fig. 8A).

### FGP: the main protein of L-EVs

FGP was identified as the major protein in L-EVs and also contained in the PM fraction (Figs. 4 and 5; Supplementary Tables S1–S3). The FG-GAP motif of FGP is conserved in extracellular proteins such as adhesins and integrins, which play roles in cell adhesion and biofilm formation (Velling et al. 1999; Absalon et al. 2011). In *Vibrio cholerae*, a Gram-negative bacterium, the Bap1 protein containing the FG-GAP motif, is involved in biofilm formation and is localized in EVs, although not as a major protein (Duperthuy et al. 2013). Thus, it is implied that FGP is localized at the outer surface of OM involved in cell adhesion and biofilm formation in *L. boryana*. Interestingly, PCC6803 does not have FGP, and FGP is largely distributed only among some filamentous cyanobacterial species such as *Anabaena* sp. PCC 7120, *Phormidium* and *Pseudoanabaena*. Thus, the presence of FGP as the major protein is one of the unique features of EVs of *L. boryana*.

### Porin isoforms in EVs

In this study, four porin isoforms were identified in EVs from WT *L. boryana* cells. Sixteen genes for porin isoforms were encoded in the genome of *L. boryana*, and the four porin isoforms detected in WT EVs corresponded to those detected in EVs of ΔDPOR grown under dark heterotrophic conditions (Usui et al. 2022). The two EVs (L-EVs and H-EVs) from WT showed almost the same protein compositions as the two EVs (Fractions 1 and 2) in ΔDPOR, respectively (Usui et al. 2022). In other words, Fractions 1 and 2 in ΔDPOR are corresponded to WT L-EVs and H-EVs, respectively. This suggested that EV formation containing large amounts of MV-Pchlide found in ΔDPOR is not a special process caused by the abnormal MV-Pchlide accumulation, but rather that the ΔDPOR cells use EVs (H-EVs) originally produced by WT cells to exclude MV-Pchlide accumulation in cells (Fig. 8B).

### Carotenoids in EVs

L-EVs contained two carotenoids, myxol glycoside and zeaxanthin, as the major pigments (Fig. 6). They are also the major pigments in the PM fraction, consistent with the idea that EVs, in general, originate from PM. EVs from PCC6803 were recovered as orange precipitates by ultracentrifugation (Biller et al. 2022b). Zeaxanthin and carotenes are also identified as the major pigments in EVs from the marine cyanobacterium *Prochlorococcus* (Biller et al. 2022a). These characteristics suggested that carotenoids, especially zeaxanthin, are the major common pigments in cyanobacterial EVs.

The increased zeaxanthin content in L-EVs from cells grown under dark heterotrophic conditions may also reflect the increased intracellular zeaxanthin content. Zeaxanthin, a component of the xanthophyll cycle, is a nonphotochemical quenching mechanism that dissipates excitation energy from Chl under excess light exposure (Eskling et al. 1997). Considering that xanthophylls, including zeaxanthin, act as the protection mechanism against oxidative stress in cyanobacteria (Zhu et al. 2010), the increase in zeaxanthin may be caused by oxidative stress under the dark heterotrophic conditions. However, the detailed mechanism behind zeaxanthin accumulation in the dark remains unclear.

### Chl biosynthetic intermediates in EVs

In this study, the Chl biosynthetic intermediates in EVs from the WT cells were analyzed in detail. EVs prepared from cells grown under different culture conditions (light and dark) were analyzed by fluorescence detection, and Chl intermediates and their derivatives were detected only in L-EVs from both cells grown under light and dark conditions (Fig. 7). Chl was not detected at all in L-EVs, and only intermediates and their derivatives before phytol attachment were specifically detected, consistent with that no Chl was detected in PM. Interestingly, three demetallated pigments, PPN, ΔMgMPE, and Pheobide, were commonly detected in both L-EVs from light- and dark-grown cells, whereas MV-Pchlide and MV-Protopheobide were additionally detected in L-EVs from dark-grown cells. A small amount of MV-Pchlide was detected in TM and PM fractions from dark-grown cells, suggesting that MV-Pchlide in the PM fraction is loaded into L-EVs. In cyanobacterial Chl biosynthesis in the dark, LPOR, which requires light for reaction, did not work, and the activity of DPOR was partially inhibited by oxygen due to the aerobic environment, resulting in a slight accumulation of Pchlide.

### Accumulation of Pheobide

Pheobide could be produced in the PSII repair cycle operating continuously in the light conditions from pheophytin in the PSII reaction centers using pheophytin dephytylase in TM (Takatani et al. 2022). Pheobide in the PM fraction from light-grown cells may be derived from the PSII repair process, although the transport process from TM to PM fraction is unknown. L-EVs may function as a molecular mechanism to remove Pheobide from TM via PM to the culture medium. However, the reason why Pheobide was also detected in L-EVs from dark-grown cells grown is still unknown.

### Accumulation of PPN and ΔMgMPE

PPN and ΔMgMPE were detected in L-EVs, but not in either TM or PM in light and dark. There are four possible causes of PPN accumulation: (1) decreased activity of Mg-chelatase and/or ferrochelatase; (2) autoxidation of the substrate protoporphyrinogen IX of protoporphyrinogen IX oxidase due to decreased activity of the enzyme; (3) decreased activity of Mg-PPN methyltransferase causing Mg-PPN accumulation, followed by demetallation; and (4) protoheme accumulation due to decreased heme oxygenase activity, followed by demetallation. A possible cause of ΔMgMPE accumulation could be the decreased activity of MPE cyclase, resulting from the decreased activity of Pchlide reductase in dark-heterotrophic conditions, followed by demetallation. However, it is unknown why the MPE cyclase activity is decreased and why these demetallated pigments accumulate under photoautotrophic conditions. However, the fact that these pigments were not detected in the CL, TM and PM fractions suggested that there is a molecular mechanism by which these pigments are efficiently transported to L-EVs.

### L-EV-mediated secretion system for Chl biosynthetic intermediates

Based on the pigment analysis of EVs, we propose a novel EV-mediated secretion system for Chl biosynthetic intermediates (Fig. 8A). In this model, accumulation of small amounts of Chl biosynthetic intermediates (such as PPN, MPE, and Chlide) continuously occurs at TM of *L. boryana* cells, and the intermediates are transported from TM to IM, presumably via “convergent membrane” (Rast et al. 2019), followed by efflux to the periplasm by an unknown transporter localized in IM. Chl intermediates in the periplasm are eventually loaded into L-EVs. Chl intermediates detected in L-EVs are commonly depleted of the central metal Mg. L-EVs from dark-grown cells also showed a high ratio of MV-Protopheobide to MV-Pchlide, suggesting that Mg-depleted intermediates may be selectively loaded into L-EVs, or that the environment in the L-EVs may facilitate demetallation.

An example of a transporter that exports PPN outside of the cell could be BCRP (ABCG2) in animal cells (Jonker et al. 2005). This transporter also has the activity to transport Pheobide (Robey et al. 2004). The genome of *L. boryana* encodes a protein (LBDG_41990) that shows significant similarity (30%) to ABCG2 and it would be a candidate for the unknown transporter.

### Loading of MV-Pchlide to H-EVs in ΔDPOR

MV-Pchlide is accumulated at a very high-density in H-EVs (Fraction 2) of ΔDPOR grown under dark heterotrophic conditions (Usui et al. 2022). This is inferred from the MV-Pchlide concentration extracted from the H-EV fraction and the significant redshift (40 nm) of the Qy peak of MV-Pchlide in the absorption spectrum of H-EVs. This may indicate that abnormal accumulation of MV-Pchlide in TM causes accumulation of MV-Pchlide in high-density OM regions promoting the loading into H-EVs (Fig. 8B). Because accumulation of misfolded proteins and peptidoglycan fragments in the periplasm promotes EV formation (Schwechheimer and Kuehn 2015), MV-Pchlide accumulation in the periplasm may be a novel factor that promotes EV formation in cyanobacteria.

### EV-mediated secretion as a novel regulatory system for Chl biosynthesis

Because Chl and its intermediates are potent strong photosensitizers, the Chl biosynthetic process is strictly regulated to minimize the accumulation of intermediates and free Chl. The EV-mediated Chl intermediate secretion system may act as a novel regulatory mechanism in Chl biosynthesis. DV-Chl *a*/*b* are contained in EVs in *Prochlorococcus* (Biller et al. 2022a), and mutants in which genes for bacteriochlorophyll biosynthesis are disrupted excrete the biosynthetic intermediates accumulating in cells to the culture medium in photosynthetic bacteria (Bollivar and Bauer 1992). Unlike plants with an efficient degradation pathway for Chls, photosynthetic prokaryotes such as cyanobacteria do not have such a Chl degradation system (Obata et al. 2019). Thus, EV-mediated efflux of Chl intermediates may play an important role as a novel regulatory mechanism for Chl biosynthesis contributing to alleviating the accumulation of Chl intermediates in cells to minimize their intracellular concentrations.

## Materials and Methods

### Cyanobacterial strains and growth condition

The filamentous cyanobacterium *L. boryana* IAM M-101 strain dg5 (Fujita et al. 1996; Hiraide et al. 2015) was used as the WT. *L. boryana* dg5 was cultivated in BG-11 medium supplemented with 20 mM HEPES-KOH (pH 7.5) and solidified with Bacto^TM^ Agar (Becton, Dickinson and Company, Franklin Lakes, NJ, USA) with a final concentration of 1.5% (w/v). Agar plates inoculated with dg5 were incubated under continuous light conditions (20 µmol m^−2^ s^−1^, fluorescent lamp, FL15EX-N-A, Hitachi, Tokyo, Japan) for 7 days at 30°C. This culture condition was used as a preculture for liquid cultivation under photoautotrophic conditions.

The pH of the BG-11 liquid medium was adjusted to 8.2 with (20 mM HEPES-KOH) for higher reproducibility of EV production under photoautotrophic conditions. BG-11 liquid medium (50 ml) was used to suspend the cells collected from the pre-culture and adjusted to an initial OD_730_ of 0.1 and bubbled with air containing 2% (v/v) CO_2_ at 30°C under continuous light for 6 days (50 µmol m^−2^ s^−1^; model SCS-2L-LED (W) Microalgae culture lighting unit, Nippon Medical and Chemical Instruments, Osaka, Japan).

For liquid cultures under dark-heterotrophic conditions, dg5 was grown on BG-11 agar plates (pH 8.2) supplemented with glucose (30 mM; BG-11+Glc), under continuous light (10 µmol m^−2^ s^−1^, fluorescent lamp FL 15EX-N-A) at 30°C for 7 days. For the main liquid culture, the cells grown on the agar plates were suspended in 100 ml BG-11+Glc liquid medium (a 500 ml Sakaguchi flask) at an OD_730_ of 0.5 and incubated at 30°C in the dark with reciprocal shaking (106 rpm; Bio-Shaker BR-3000LF, TITEC, Koshigaya, Japan) for 5 days.

### Preparation of EV fractions

EVs were prepared described previously (Usui et al. 2022). The culture supernatant was collected by centrifugation (6,000 rpm, 10 min, 4°C; R10A2 rotor, Himac CR21F; Eppendorf Himac Technologies, Hitachinaka, Japan). For cultures grown under photoautotrophic conditions, the supernatant was filtered through a 0.22 µm filter (150 ml vacuum filter system; Corning, New York, USA) to remove cells and debris. For cultures grown under dark-heterotrophic conditions because the culture medium tended to clog a 0.22 µm filter, the culture supernatant of the culture was prepared by centrifugation (6,000 rpm, 10 min, 4°C; R10A2 rotor, Himac CR21F). The crude EV fractions were prepared by ultracentrifugation (45,000 rpm, 1 h, 4°C; S50A rotor, Himac Ultracentrifuge CS100GXII; Eppendorf Himac Technologies) of the thus prepared supernatants.

The resulting precipitate was suspended in 1 ml phosphate-buffered saline (PBS) containing 1 mM EDTA (all buffer solutions referred to as PBS hereafter contain 1 mM EDTA). For EVs from cultures grown under photoautotrophic conditions, this suspension was used as the crude EV fraction. For EVs from cultures grown dark heterotrophic conditions, the precipitate obtained by ultracentrifugation was suspended and centrifuged again at low-speed (6,000 rpm, 2 min, 4°C; ARO 15-24 rotor, centrifuge MX-301; Tommy Seiko, Tokyo, Japan) to completely remove any mixed cells, and the supernatant was used as the crude EV fraction. Stepwise sucrose density gradient centrifugation for two EV fractions was carried out as described previously (Usui et al. 2022).

### Preparation of membrane fractions

The preparation of membrane fractions is shown in Supplementary Figure S1. Cells grown under photoautotrophic and dark heterotrophic conditions were collected using centrifugation (6,000 rpm, 10 min, 4°C; R10A2 rotor, Himac CR21F, Eppendorf Himac Technologies) and washed with a small amount of PBS. After washing, the suspended cells were dispensed into 1.5 ml tubes at volumes of 500 µl. Approximately 0.1 g of glass beads (150–212 µm; Sigma-Aldrich, St. Louis, MO, USA) were added to the samples, and cells were disrupted using a bead beater (an LA intensity, 1 h, 4°C; BugCrasher GM-01; TITEC, Koshigaya, Japan). To remove cell debris and glass beads, samples were centrifuged (3,000 rpm, 3 min, 4°C; ARO 15–24 rotor, MX-301), and phenylmethylsulfonyl fluoride (PMSF) was added to the supernatants to a final concentration of 0.2 mM. The resulting CLs were stored at –80°C.

Because TM, PM, and CW fractionation from cyanobacteria was performed by sucrose density gradient centrifugation (Murata and Omata 1988; Kowata et al. 2017; Supplementary Fig. S1). First, CL was ultracentrifuged (19,000 rpm, 15 min, 4°C; S50A rotor, Himac CS100GXII) to obtain a precipitate of high-density membrane fraction (Supplementary Fig. S1A, (2)). The supernatant was further ultracentrifuged (40,000 rpm, 30 min, 4°C; S50A rotor; Himac CS100GXII) to obtain precipitates of the low-density membrane fraction (Supplementary Fig. S1 A (1)). These two precipitates were suspended in small volumes of PBS, and aliquots (1 ml) of the precipitates were subjected to stepwise sucrose density gradient centrifugation (2.5, 1.6, and 0.6 M; 25,000 rpm, 20 h, 4°C; SRP28SA rotor, Himac CP65β, Eppendorf Himac Technologies; Supplementary Fig. S1B, C, E, and F). From the low-density membrane fraction, the orange band at the 0.6 M to 1.6 M interface was collected as the crude PM fraction, and three to five times the volume of PBS was added to wash this fraction. To remove other membrane fractions, ultracentrifugation (19,000 rpm, 10 min, 4°C; S50A rotor; Himac CS100GXII) was performed and the supernatant was ultracentrifuged again (40,000 rpm, 1 h, 4°C; S50A rotor; Himac CS100GXII). The resulting precipitate was suspended in PBS, the final PM fraction. For the high-density membrane fractions, green and orange bands were collected as TM and CW fractions, respectively. After each fraction was collected, three to five times the volume of PBS was added, and the precipitate obtained by ultracentrifugation (27,000 rpm, 30 min, 4°C; S50A rotor; Himac CS100GXII) was again suspended in a small volume of PBS. The same sucrose density gradient centrifugation was repeated to prepare the TM and CW fractions (Supplementary Fig. S1D and G). Each membrane fraction was stored at –80°C in the dark.

### Determination of protein concentration

Protein concentrations were determined using the BCA method (TaKaRa BCA Protein Assay Kit; Takara, Shiga, Japan) after 1% SDS was added to solubilize the samples. Bovine serum albumin was used as the standard.

### Pigment analysis

Methanol was added to each sample (final concentration 90%, v/v)). The samples were suspended well and kept on ice for 20 min in the dark, followed by centrifugation at 15,000 rpm at 4°C for 10 min (ARO15-24 rotor, MX-301). Subsequently, an aliquot (20 µl) of the supernatant was analyzed using HPLC (Zapata et al. 2000) with a column (4.6 × 150 mm Symmetry C8 3.5 µm; Waters, Milford, MA, USA) coupled with a Shimadzu LC series HPLC system (Shimadzu, Kyoto, Japan). Pigments were detected by absorption at 440 nm (SPD-20AV; Shimadzu), and the absorption spectrum of the eluted pigments was continuously monitored by a photodiode array detector (SPD-M20A; Shimadzu). The fluorescence of Chl intermediates and their derivatives was detected using a fluorescence detector (RF-20AXS; Shimadzu). Three sets of different fluorescence emission and excitation wavelengths were employed for the pigment detection: λ_em_/λ_ex_ 660/430, 630/400, and 600/417 nm.

### Preparation of Chlide and Pheobide

Chlide was prepared from a Δ*bchZ/bchF* mutant of *Rhodobacter capsulatus* (CB1200; Bollivar et al. 1994) as described previously described (Nomata et al. 2006). Tween 20 (final concentration 0.2%, v/v) was added to the culture (RCV-2/3PY; Yang et al. 1989) to promote pigment production. The prepared Chlide was dissolved in dimethyl sulfoxide (DMSO). The final Chlide sample was prepared by diluting of 5 µl Chlide in DMSO with 95 µl of water. To prepare Pheobide, the central Mg ion was removed by acid treatment with a drop of hydrochloric acid (35–37%) to the final Chlide sample.

### Preparation of MPE and demetallated MPE

MPE and ΔMgMPE were prepared from a mutant Δ*operon* of PCC6803 (Aoki et al. 2014). The Δ*operon* mutant was isolated by replacing of the *chlA_II_-ho2-hemN* operon with the kanamycin resistance cartridge (Aoki et al. 2014). Δ*operon* cannot grow under microoxic conditions due to the removal of the three genes essential for Chl and heme biosynthesis under low-oxygen conditions. When Δ*operon* cells were exposed to microoxic conditions, the cells accumulated various Chl biosynthetic intermediates; the main pigments were MPE, ΔMgMPE and PPN (Supplementary Fig. S6). First, Δ*operon* was cultivated on a BG-11 agar plate containing kanamycin (15 µg ml^−1^) under continuous light (20 µmol m^−2^ s^−1^, fluorescent lamp, FL15EX-N-A) for 7 days at 30°C. The agar plate was transferred to an anaerobic jar (AnaeroPac square jar; Mitsubishi Gas Chemical, Tokyo, Japan) containing an oxygen removal pouch (AnaeroPac Kenki, Mitsubishi Gas Chemical) for 5 days. The maintenance of anaerobic conditions was confirmed by a strip of methylene blue-containing indicator (Dry Anaerobic Indicator; BD BBL, Franklin Lake, NJ, USA) in the jar. After anaerobic incubation, cells were suspended in 100 µl water and used for HPLC analysis as described above.

### SDS-PAGE and Western blot analysis

SDS-PAGE and Western blot analysis were performed as described previously (Usui et al. 2022). The anti-PsbA antiserum (Agrisera, Vännäs, Sweden) was used at a ratio of 1:1,000.

### MS analysis

MS analysis was carried out essentially as described previously (Usui et al. 2022) using MS (QTRA5500, Sciex, Tokyo, Japan) and nanoLC chromatography (Eksigent LC425, Sciex) with 0.1% formic acid, 2–35% acetonitorile. An aliquot (8 µL) of the sample was injected into a nano HPLC capillary column (NTCC-360/75-3-125 Nikkyo Technos, Tokyo, Japan) and inserted into the MS. For qualitative analysis, information-dependent acquisition was performed with an MS scan range of 400 to 1,000 Da and an enhanced product ion scan range of 100 to 1,000 Da. Results were analyzed with ProteinPilot^TM^ Software 5.0 (Sciex), and those with a total ProtScore of> 10 are listed in each table (Supplementary Tables S1–S3).

### TEM

In negative staining, a small aliquot (5 µl) of the purified EV fraction was placed on a copper grid (200 mesh; Nissin EM, Tokyo, Japan) coated with formvar (Nissin EM) and incubated for 5 min. The excess solution was absorbed by filter paper, and 4 µl of 2% uranium acetate solution were placed on the grid and stained for 2 min. After staining, the excess staining solution was absorbed by filter paper and dried for a few minutes. The stained grids were observed using TEM (H-7500, Hitachi), and images were taken using a CCD camera (Advanced Microscopy Technique, Woburn, MA, USA).

## Data Availability

Data are contained within the article or Supplementary Materials. All raw data have been deposited into the data management system of Nagoya University.

## Acknowledgments

We thank Takao Oi and Mitsutaka Taniguchi (the Laboratory of Plant Physiology and Morphology, Graduate School of Bioagricultural Sciences, Nagoya University) for supports of TEM observation. We thank Takahiro Arase for preparation of Chlide, and Asako Segawa for technical help. Takafumi Yamashino, Mari Banba and all members of the Laboratory of Molecular and Functional Genomics for discussion and technical help.

## Author contributions

K.U. and Y.F. conceived the study and designed the experiments. K.U. performed all experiments. K.U. conducted TEM observation. H.M. and K.U. conducted MS analysis. K.U. and Y.F. wrote the manuscript. All authors reviewed the manuscript. All authors have read and agreed to the published version of the manuscript.

