## Supplementary materials for "Extracellular vesicle-mediated secretion of chlorophyll biosynthetic intermediates in the cyanobacterium *Leptolyngbya boryana*"

**Supplementary Figures: 7**

**Supplementary Figure S1.**	Purification of each membrane fraction

**Supplementary Figure S2.**	PDA absorption spectra of carotenoids

**Supplementary Figure S3.**	PDA absorption spectrum of MV-Protopheobide

**Supplementary Figure S4.**	HPLC profiles and PDA absorption spectra of Chlide and Pheobide

**Supplementary Figure S5.**	PDA absorption spectra of pigments that appeared to be a Chlide derivative

**Supplementary Figure S6.**	HPLC profiles of PPN, MPE, and Mg-demetallated MPE

**Supplementary Figure S7.**	HPLC analysis to detect MPE derivatives

**Supplementary Tables: 3**

**Supplementary Table S1.**	MS of the major proteins in the EV fractions

**Supplementary Table S2.**	MS of the major proteins in PM and CW fractions of WT grown under photoautotroph conditions

**Supplementary Table S3.**	MS of the major proteins in PM and CW fractions of WT grown under dark heterotrophic conditions

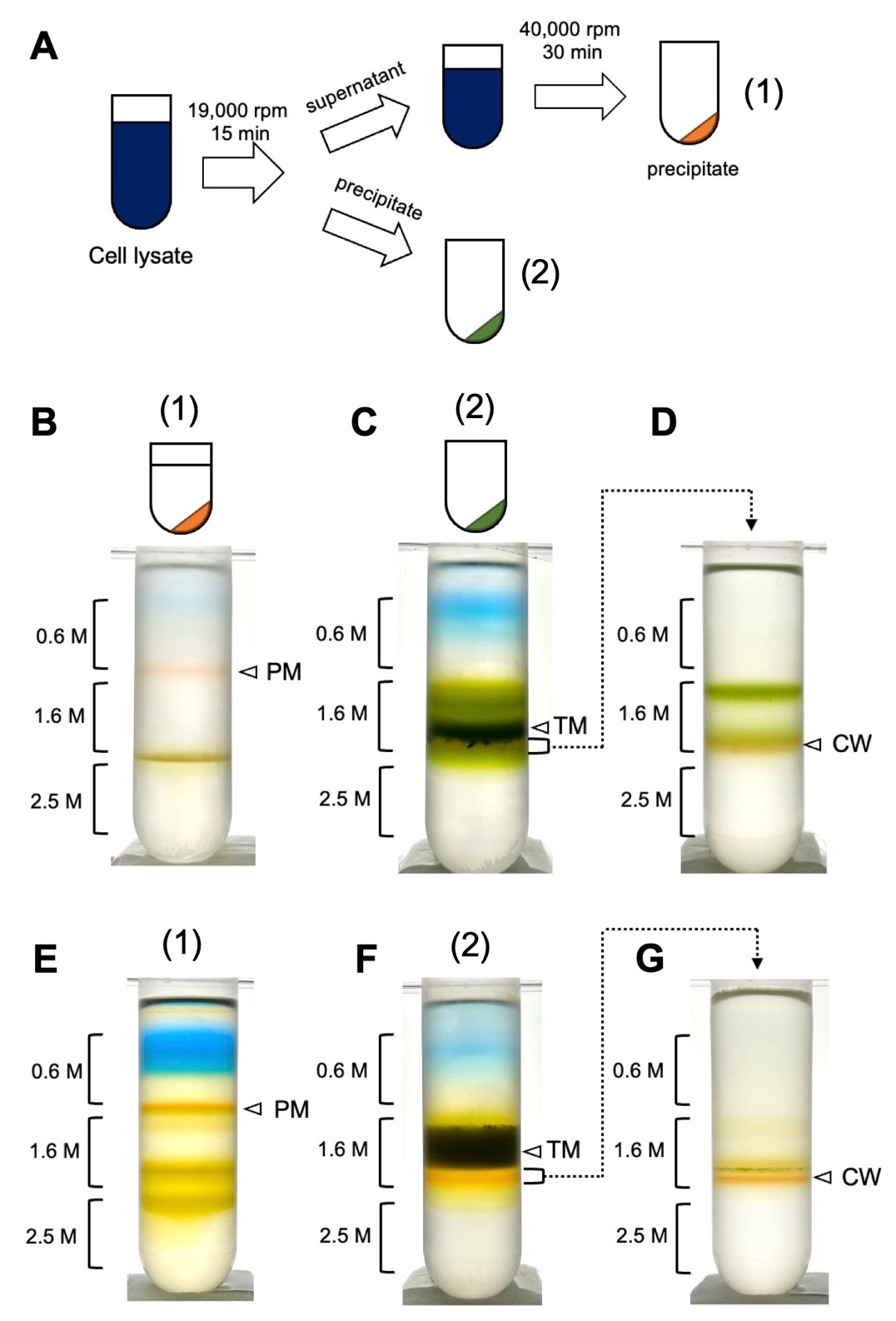

**Supplementary Fig. S1.** Purification of each membrane fraction. **A.** Differential ultracentrifugation was employed to extract high-density and low-density membrane fractions from CL. **B–G.** Sucrose density gradient centrifugation of the membrane fractions of cells grown under both light (**B**–**D**) and dark (**E–G**) conditions. Sucrose density gradient centrifugation of the high-speed precipitate (a); the orange band at the 0.6 M to1.6 M interface was collected as the PM fraction (**B** and **E)**. Sucrose density gradient centrifugation of the low-speed precipitate (b); the dark green band slightly above the 1.6 M to 2.5 M boundary was obtained as the TM fraction (**C** and **F**). The 1.6 M-2.5 M boundary underwent density gradient centrifugation again (**C** and **F**). The orange band at the 1.6 M to 2.5 M boundary was collected as the CW fraction (**D** and **G**).

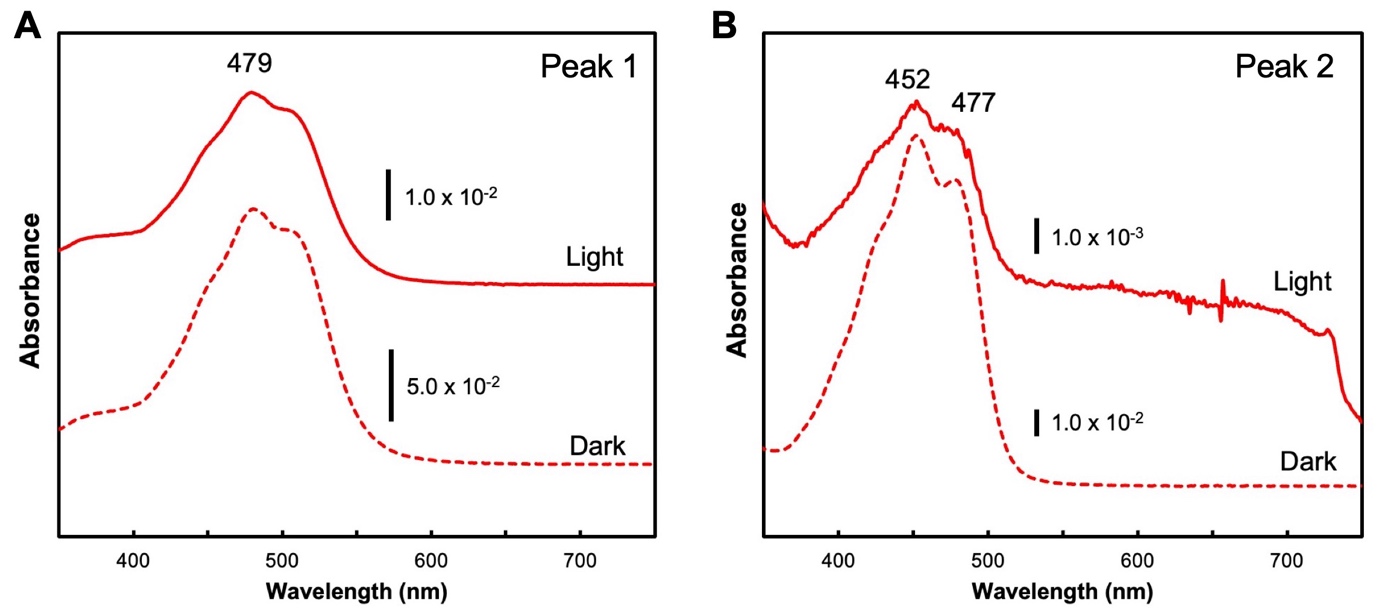

**Supplementary Fig. S2.** PDA absorption spectra of carotenoids. PDA absorption spectra corresponding to peak 1 (**A**; myxol glycoside) and peak 2 (**B**; zeaxanthin) in HPLC profiles of L-EV fractions in Fig. 6. Solid and dashed lines are from HPLC profiles of L-HVs from cells grown under photosynthetic and dark heterotrophic conditions, respectively.

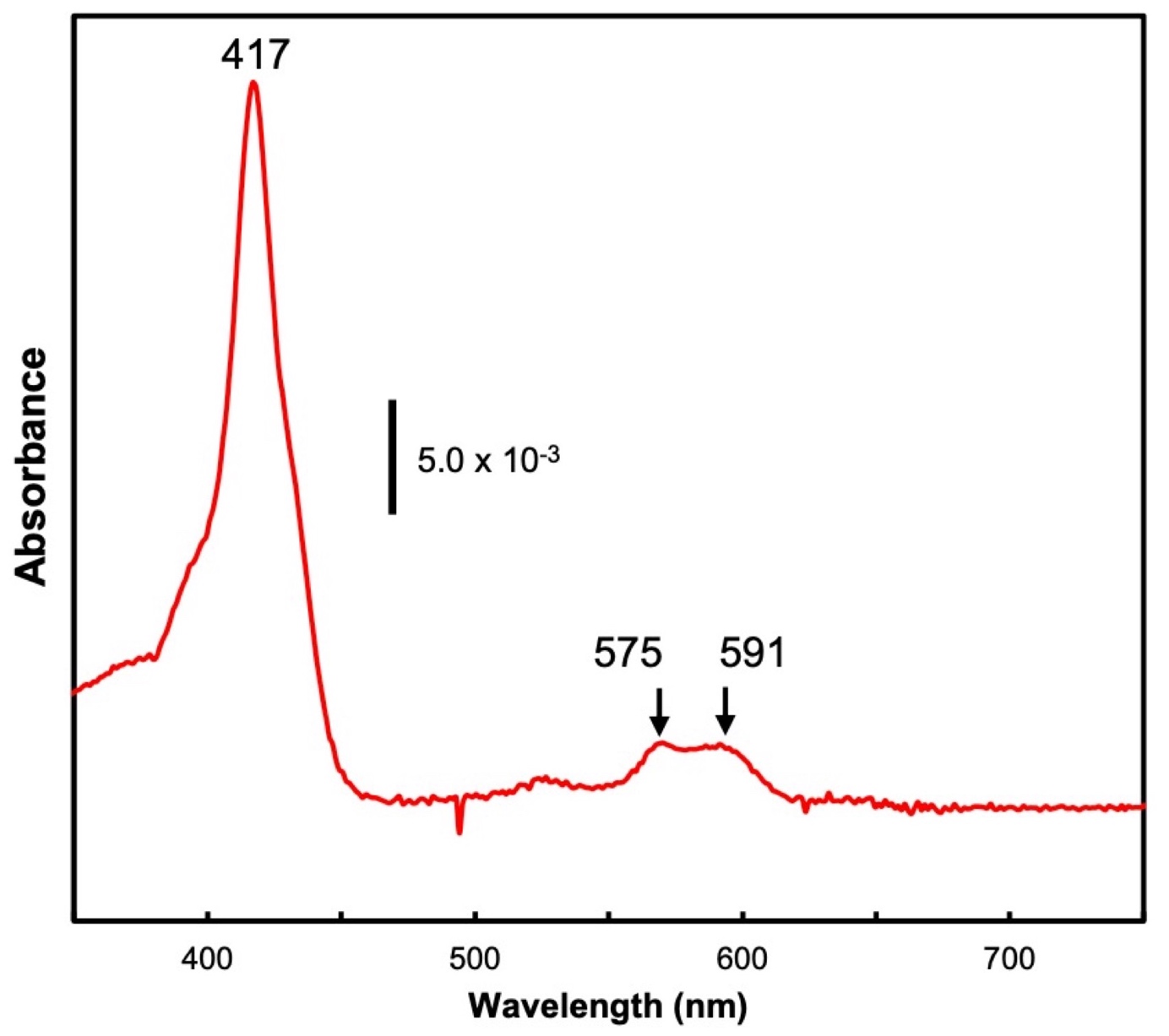

**Supplementary Fig. S3.** PDA absorption spectrum of MV-Protopheobide. PDA absorption spectrum of the pigment (MV-Protopheobide) eluted at ~10.0 min (Fig. 6B, peak 5) in the L-EV fraction from dark-grown cells.

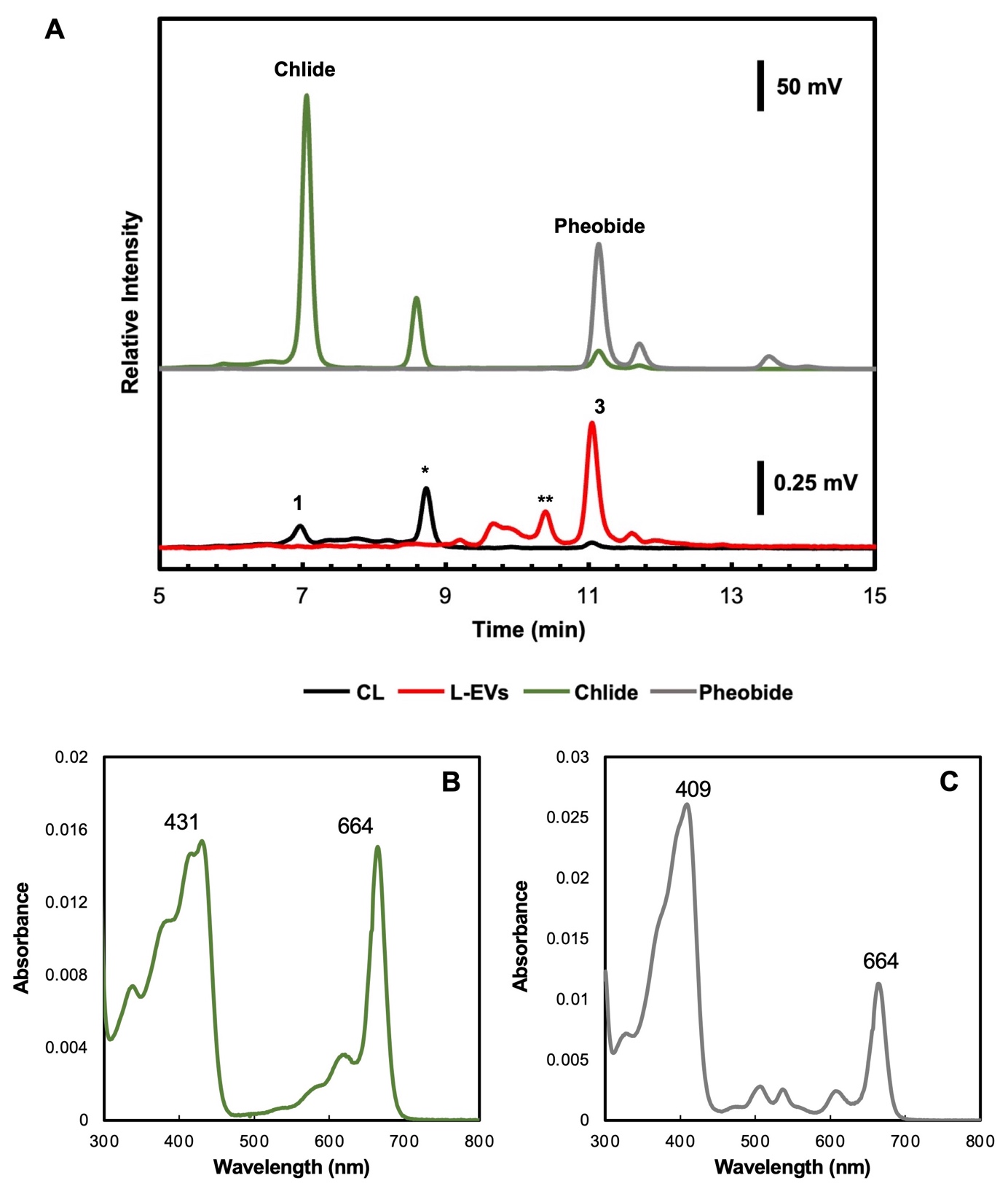

**Supplementary Fig. S4.** HPLC profiles and PDA absorption spectra of Chlide and Pheobide.	**A.** HPLC profiles of Chlide (dark green trace) purified from the *∆bchZ/bchF* mutant (CB1200) of the photosynthetic bacterium *R. capsulatus* and Pheobide (gray trace) prepared by acid treatment of the purified Chlide sample (fluorescence detection; λ_em_/λ_ex_ 660/430 nm). Elution profiles of Chlide and Pheobide were compared to those of CL (black trace) and L-EVs (red trace) from light-grown cells. Peak numbers in CL and L-EVs are the same as in Fig. 7. The asterisk (*) indicates a Chlide derivative (Supplementary Fig. S5A). **B and C.** PDA absorption spectra of Chlide (**B**; peak 1) and Pheobide (**C**; peak 3) are also shown.

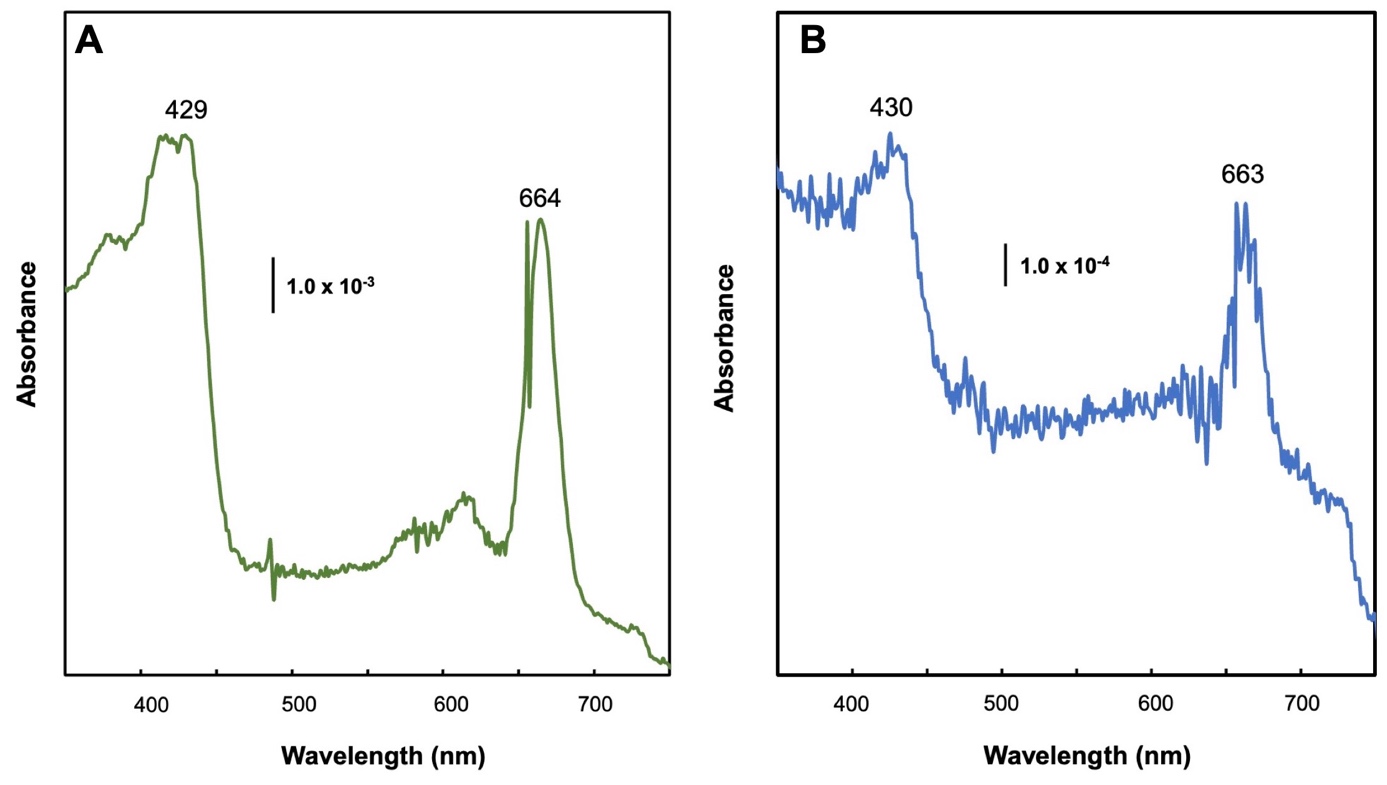

**Supplementary Fig. S5.** PDA absorption spectra of pigments that appeared to be a Chlide derivative.	**A.** PDA absorption spectrum of pigment that appears to be a Chlide derivative, eluted at 8.5 min (peak 2) in Supplementary Fig. S4A. **B.** PDA absorption spectrum of the pigment eluted at 8.7 min elution time in the TM fraction of Fig. 7A (peak *).

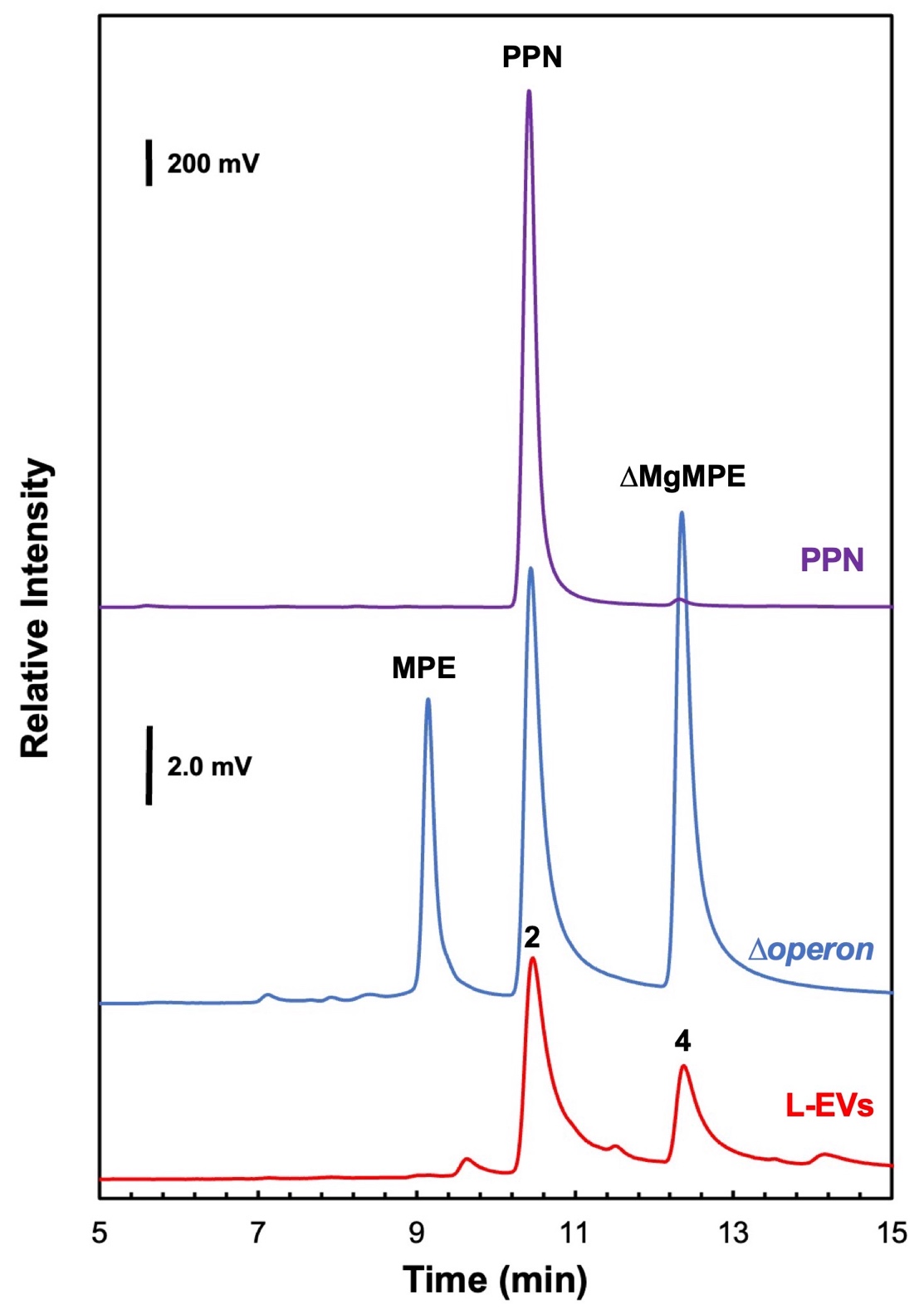

**Supplementary Fig. S6.** HPLC profiles of PPN, MPE, and Mg-demetallated MPE.	HPLC profile (fluorescence detection; λ_em_/λ_ex_ 630/400 nm) of standard samples for PPN (purple trace), MPE and ∆MgMPE (blue trace), and L-EVs (red trace) from cells grown under light conditions. Standard samples of MPE and ∆MgMPE were prepared by extracting the pigment from *Synechocystis* sp. PCC 6803 *∆operon* cells grown under anaerobic conditions for 5 days with 90% methanol (Aoki et al. 2014). The peak numbers in L-EVs correspond to those in Fig. 7A.

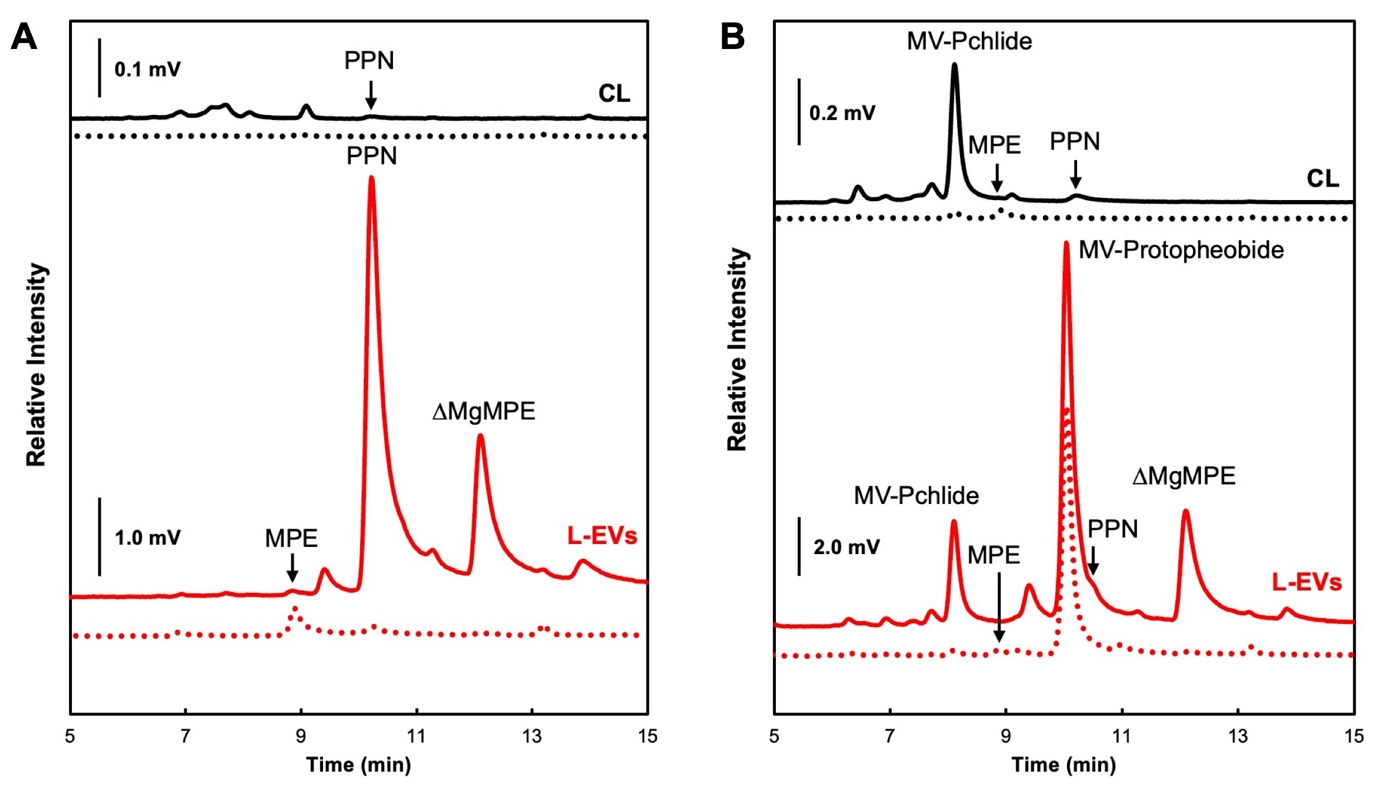

**Supplementary Fig. S7.** HPLC analysis to detect MPE derivatives.	HPLC profiles of pigments extracted from CL (black trace) and L-EVs (red trace) of the light- and dark-grown cells are shown in **A** and **B**, respectively. Solid and dotted traces are the profiles at λ_em_/λ_ex_ 630/400 and 600/417 nm, respectively, used for specific detection of MPE derivatives. The protein concentration in each fraction was adjusted to 0.1 mg/ml, and an aliquot (20 µl) of each extracted pigment was injected.

**Supplementary Tables**

**Supplementary Table S1.** MS of the major proteins in the EV fractions.

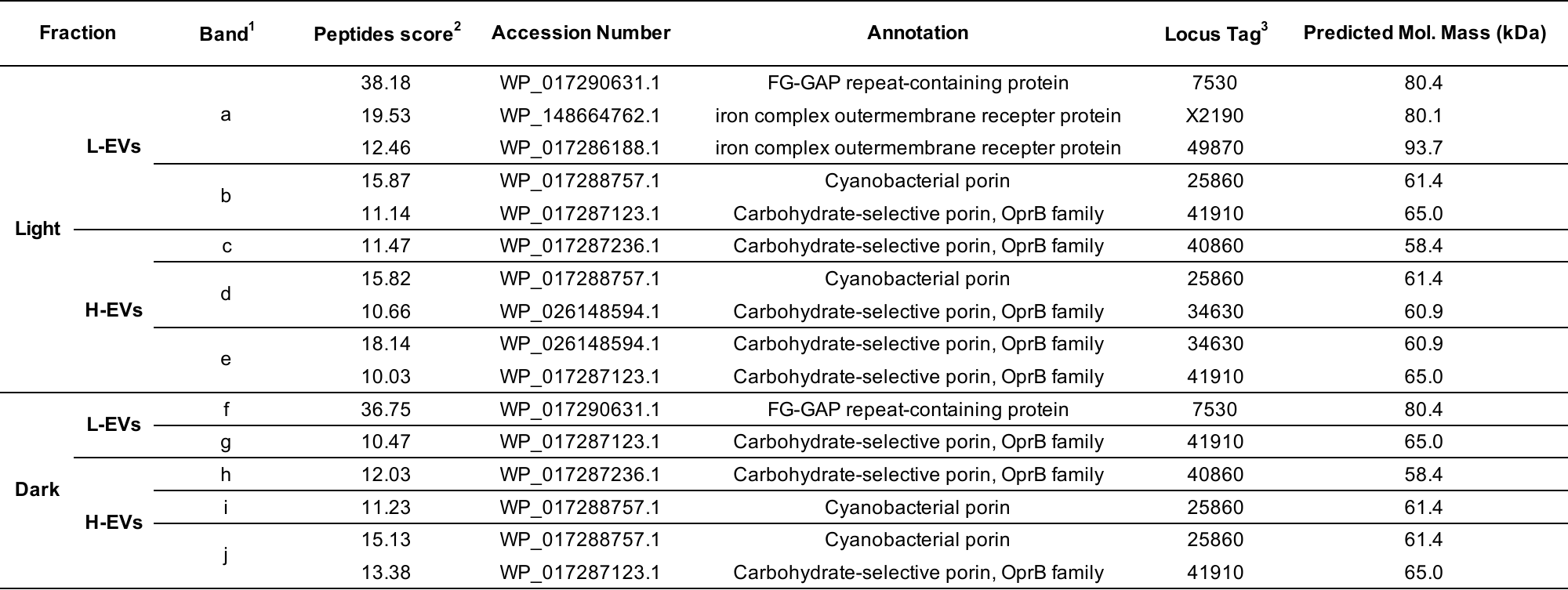

^1^ Bands a–j (Fig. 4) were excised and analyzed by MS. ^2^ ProtScore. ^3^ LBDG_xxxxx.

**Supplementary Table S2.** MS of the major proteins in PM and CW fractions of WT grown under photoautotroph conditions.

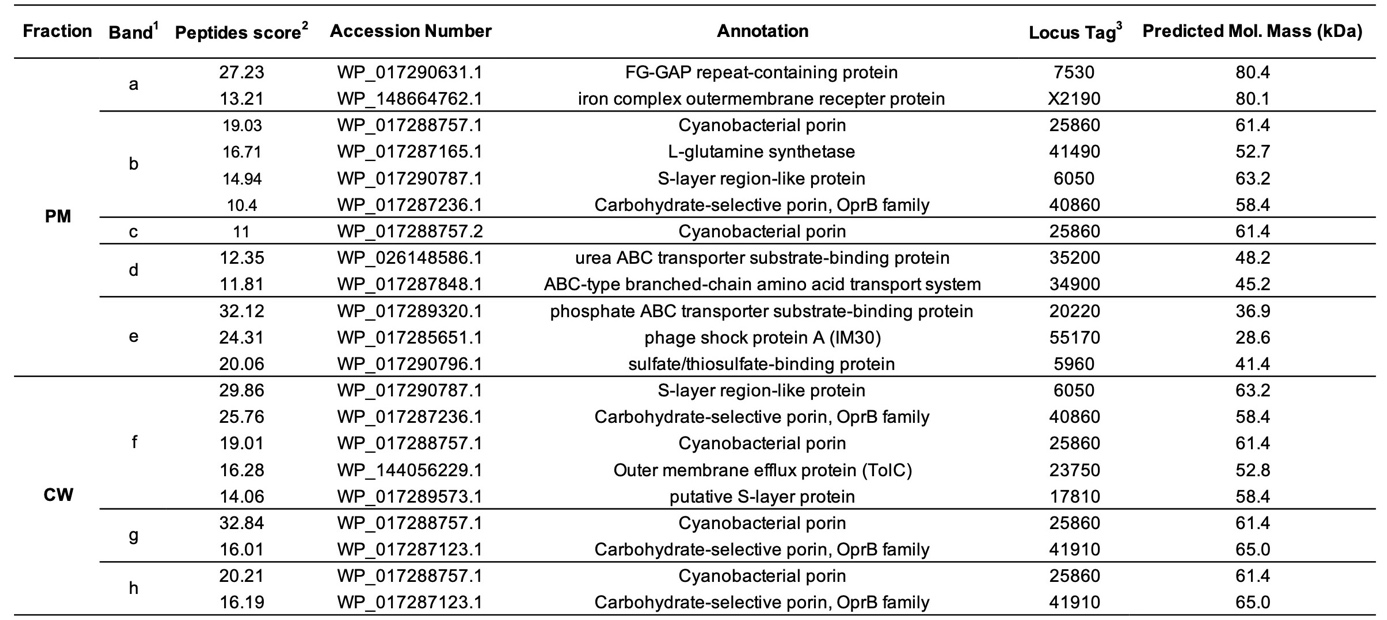

^1^ Bands a–h in Fig. 5A were excised and analyzed by MS. ^2^ ProtScore. ^3^ LBDG_xxxxx

**Supplementary Table S3.** MS of the major proteins in PM and CW fractions of WT grown under dark heterotrophic conditions.

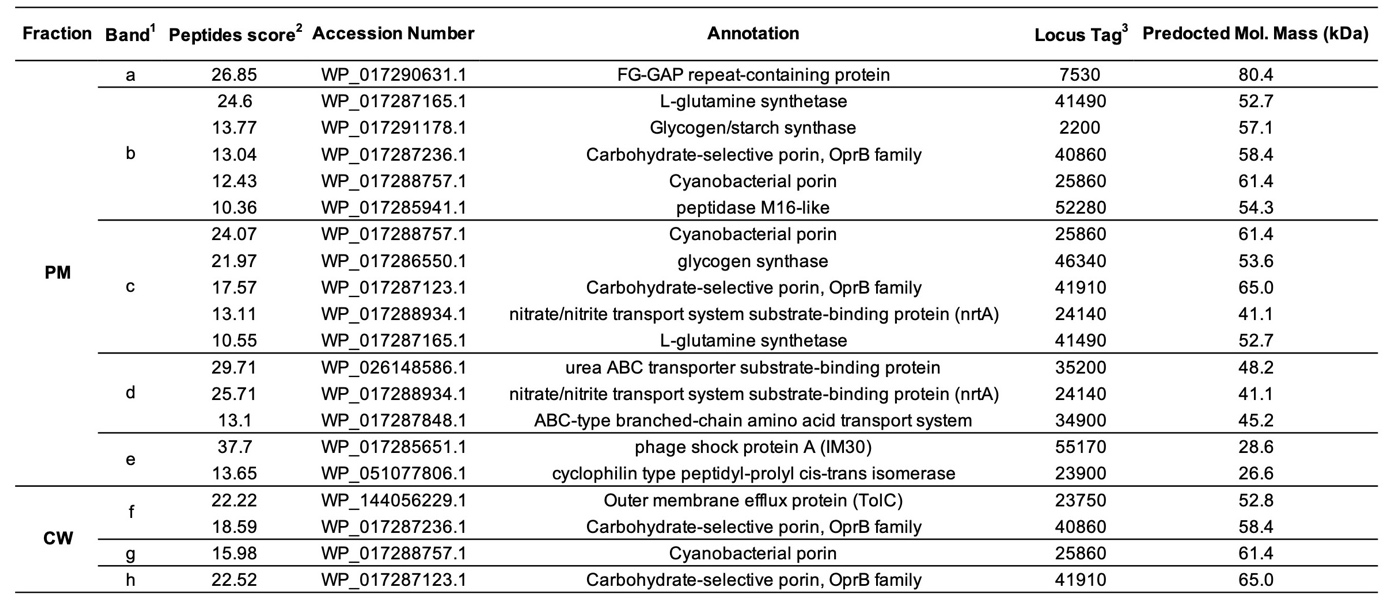

^1^ Bands a–h in Fig. 5B were excised and analyzed by MS. ^2^ ProtScore. ^3^ LBDG_xxxxx
